# Messenger RNA cap methylation by PCIF1 attenuates the interferon-β induced antiviral state

**DOI:** 10.1101/2020.12.17.423296

**Authors:** Michael A. Tartell, Konstantinos Boulias, Gabriela Brunsting Hoffmann, Eric Lieberman Greer, Sean P. J. Whelan

## Abstract

Interferons induce cell intrinsic responses associated with resistance to viral infection. To overcome the suppressive action of interferons and their downstream effectors viruses have evolved diverse mechanisms. Working with vesicular stomatitis virus (VSV) we report a role for the host cell N6-adenosine mRNA cap-methylase, phosphorylated C-terminal domain interacting factor 1 (PCIF1), in attenuating the antiviral activity of interferon-β. Using cell based and *in vitro* biochemical assays we demonstrate that PCIF1 efficiently modifies VSV mRNA cap structures to m^7^Gpppm^6^A_m_, and we identify the *cis*-acting elements required for this modification. Under basal conditions, N6-methylation of VSV mRNA cap structures is functionally inert with regard to mRNA stability, translation and viral infectivity. Induction of an antiviral state by treatment of cells with interferon-β prior to infection uncovered a functional role for PCIF1 in attenuation of the antiviral response. Cells lacking PCIF1 or expressing a catalytically inactive PCIF1, exhibit an augmented effect of interferon-β in the inhibition of viral replication and gene expression. This work identifies a function of PCIF1 and cap-proximal m^6^A_m_ in attenuation of the host response to VSV infection that likely extends to other viruses.

**Significance:** The cap structure present at the 5’ end of eukaryotic mRNAs regulates RNA stability, translation, and marks mRNA as self, thereby impeding recognition by the innate immune system. Cellular transcripts beginning with adenosine are additionally modified at the N6 position of the 2’-O methylated cap-proximal residue by the methyltransferase PCIF1 to m^7^Gpppm^6^A_m_. We define a function for this N6-adenosine methylation in attenuating the interferon-β mediated suppression of viral infection. Cells lacking PCIF1, or defective in its enzymatic activity, augment the cell intrinsic suppressive effect of interferon-β treatment on vesicular stomatitis virus gene expression. VSV mRNAs are efficiently methylated by PCIF1, suggesting this contributes to viral evasion of innate immune suppression.

## Introduction

Eukaryotic messenger RNAs possess a 5’ cap structure that functions in their stability, translation, and helps discriminates host from aberrant RNA by the innate immune system (1–4). That mRNA cap structure is formed by the actions of an RNA triphosphatase that converts pppRNA to ppRNA which serves as substrate for an RNA guanylyltransferase to transfer GMP derived from GTP onto the 5’ end of the RNA to yield GpppRNA (1, 3, 5). Methylation of that 5’ cap-structure by a guanine-N-7 methylase yields m^7^GpppRNA, which is modified by a ribose-2’-O methylase to yield m^7^GpppN_m_ (1, 3). Known activators of the innate immune system include triphosphate RNA which is recognized by the host pattern recognition receptor, retinoic acid inducible gene-1 (RIG-1) (4, 6), and cap-structures that lack ribose-2’-O methylation which renders translation of those RNAs susceptible to inhibition by interferon-induced protein with tetratricopeptide repeats 1 (IFIT1) (4, 7).

Internal RNA modifications also have important functional consequences for the fate of mRNA, among which is N6-methyladenosine (m^6^A). The methyltransferase complex METTL3/METTL14 is responsible for m^6^A methylation, which regulates diverse functions in mRNA localization, stability, splicing, and translation (8, 9). The RNA modification N6, 2’O di-methyladenosine (m^6^A_m_) present at the cap-proximal position (m^7^Gpppm^6^A_m_) is regulated separately from m^6^A (10). Cap proximal m^6^A_m_ is present on approximately 30% of cellular mRNA (11–16), but its function is enigmatic. The host RNA polymerase II associated phosphorylated-CTD interacting factor 1 (PCIF1), catalyzes formation of cap proximal m^6^A_m_ (11–14) that is reported to increase (11, 12, 17), decrease (13), or have no consequence for (18) mRNA stability and translation.

Vesicular stomatitis virus (VSV), a non-segmented negative-sense RNA virus, replicates in the host-cell cytoplasm transcribing 5 mRNAs from the viral genome (19). The viral large polymerase protein (L) contains the enzymatic activities necessary for transcription of the 5 mRNAs, including the co-transcriptional addition of a 5’ methylated cap-structure (m^7^GpppA_m_) and synthesis of the 3’ poly-A tail (20). The polymerase synthesizes the mRNAs by recognizing conserved stop and start sequences within each gene so that each mRNA contains an identical 5’ structure m^7^GpppA_m_ACAG (21–23). VSV mRNAs isolated from cells are additionally N6-methylated at the cap-proximal A_m_ by a presumed cellular methylase to yield m^7^Gpppm^6^A_m_ACAG (24). The efficiency of VSV transcription is such that at least 65% of total cytoplasmic mRNA corresponds to the 5 VSV mRNAs by 6 hours post infection (25). The 5 VSV mRNAs and their protein products have been extensively characterized biochemically (26), and as a result provide unique probes into the function(s) of m^6^A_m_.

Here, we demonstrate that VSV mRNAs are efficiently modified at the cap-proximal nucleotide by host PCIF1. In contrast to the substrate requirements for cellular mRNA modification, the PCIF1-dependent N6-methylation of VSV mRNA is independent of prior guanine-N7-methylation of the mRNA cap. Under basal conditions, VSV mRNA stability and translation are unaffected by the presence of m^6^A_m_, and viral replication is unaltered. Activation of an antiviral response by treatment of cells with interferon-β uncovers a function for PCIF1. Cells lacking PCIF1 or expressing a catalytically inactive variant of the protein exhibit an augmented suppression of viral gene expression and infection upon interferon-β treatment. The attenuation of the antiviral activity of interferon-β was dependent upon the catalytic activity of PCIF1, thus defining a functional role of this mRNA cap methylation in evading antiviral suppression of gene expression. This work defines a role of PCIF1 dependent methylation of mRNA cap-structures in the attenuation of the antiviral response in VSV infected cells that likely extends to other viruses.

## Results

### Analysis of VSV mRNA cap-structures isolated from cells

To examine the methylation status of VSV mRNA cap structures, we infected 293T cells at a MOI of 3, labeled viral RNA by metabolic incorporation of [^32^P]-phosphoric acid in the presence of actinomycin-D from 3-7 hpi, which selectively inhibits host cell transcription, and isolated total poly(A)+ RNA. Following hydrolysis with nuclease P1 to liberate mononucleotides, and cap-clip acid pyrophosphatase to digest the mRNA cap-structure (Fig 1A), the products were resolved by two-dimensional thin-layer chromatography (2D-TLC) (27) (Fig 1B). The identity of specific spots was determined by their comigration with co-spotted chemical markers (Fig S1A) (17, 27). Analysis of RNA from uninfected cells yields low levels of products that comigrate with pA, pC, pG, and pU reflecting residual actinomycin-D resistant synthesis of RNA in the cell, but no detectable methylated nucleotides (Fig 1B). Nuclease P1 digestion of RNA from infected cells gave rise to abundant pA, pC, pG and pU and two additional spots that comigrate with markers for A_m_ and m^6^A (Fig 1B). As nuclease P1 leaves the mRNA cap-structure intact, the presence of A_m_ and m^6^A must reflect internal modifications of the viral mRNA, although our analysis cannot discriminate which positions are modified. Prior studies on VSV mRNAs report the presence of A_m_ at the second transcribed nucleotide (24), which may account for some of this internal methylation. Further hydrolysis of the RNA purified from infected cells with cap-clip pyrophosphatase leads to the appearance of an additional spot that comigrates with m^6^A_m_, and an increase in intensity of the A_m_ spot (Fig 1B). This result demonstrates that VSV mRNA cap structures contain m^7^Gpppm^6^A_m_, as m^6^A_m_ only appears upon cap-structure hydrolysis. We interpret the increase in A_m_ following cap-clip treatment as reflective of the presence of a minority of transcripts containing only A_m_ at the cap-proximal nucleotide (24). Quantitative analysis of each spot reveals over 85% of the cap-proximal nucleotides are m^6^A_m_, consistent with previous reports (24). Experiments performed in HeLa cells gave similar results, though no internal m^6^A was detectable in these cells (Fig S1B). We conclude that VSV mRNAs synthesized in 293T and Hela cells contain primarily m^7^Gpppm^6^A_m_ and m^7^Gpppm^6^A_m_A_m_ cap-structures.

**Figure 1:**
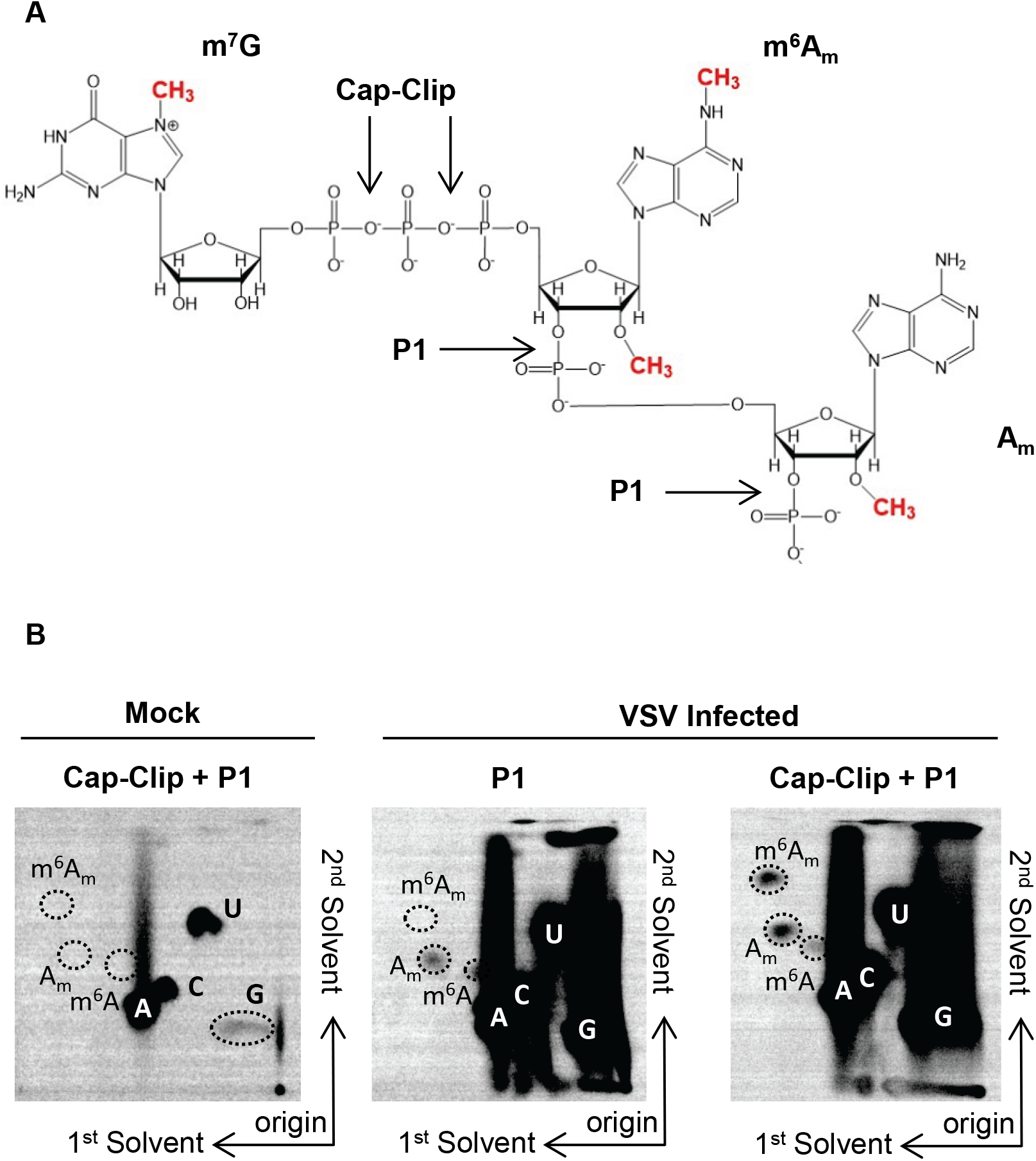
VSV mRNAs contain a 5’ m^7^Gpppm^6^A_m_ cap-structure. **(A):** VSV mRNA cap-structures present in infected cells. VSV mRNAs contain the conserved 5’ gene start sequence AACAG, including the m^7^GpppA_m_ cap structures synthesized by the VSV polymerase, N6-methylation at the cap-proximal first nucleotide, and 2’O methylation at the second nucleotide made by the cell. Sites of nuclease P1 and cap-clip pyrophosphatase cleavage are marked. **(B):** VSV contains m^6^A_m_ at the cap-proximal nucleotide. 293T cells were infected with VSV at a MOI of 3, cellular transcription halted by adding 10 μg ml^−1^ actinomycin D at 2.5 hpi, and viral RNA labeled by metabolic incorporation of 100 μCi ml^−1^ [^32^P] phosphoric acid from 3-7 hpi. Total cellular RNA was extracted and following poly(A) selection hydrolyzed by the indicated nucleases into monophosphates that were resolved by 2D-TLC and detected by phosphorimaging (representative image; n=3). Solvents run in the first and second dimensions are marked.

### PCIF1 modifies VSV mRNA

To determine whether VSV mRNA are modified by PCIF1, the cellular N6-methyltransferase for mRNA (Fig S2) (11–14), we infected *PCIF1* knockout (KO) cells and performed RNA analysis as above. Viral mRNA isolated from PCIF1 KO cells (HeLa or 293T) lacked detectable levels of m^6^A_m_ (Fig 2A, Fig S1C). Cap-proximal m^6^A_m_ was restored upon add back of PCIF1 but not a catalytically inactive mutant PCIF1_SPPG_ (Fig 2B). Methylation reactions performed *in vitro* demonstrate that purified PCIF1 but not PCIF1_SPPG_ are responsible for m^6^A_m_ on VSV mRNA (Fig 2C, S1D). Collectively these results demonstrate that PCIF1 is necessary and sufficient for formation of cap-proximal m^6^A_m_ on VSV mRNA.

**Figure 2:**
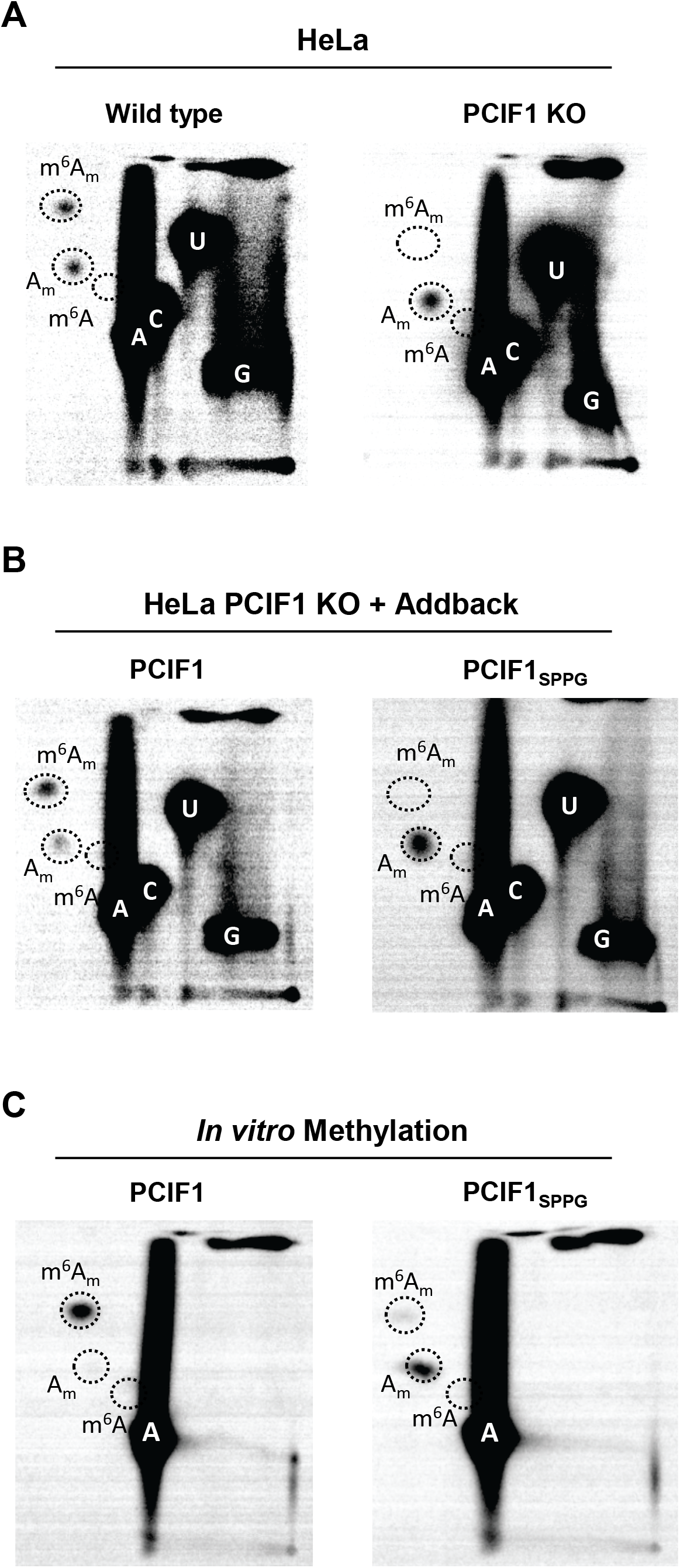
PCIF1 is the cap-proximal N6-methyltransferase. **(A):** CRISPR-mediated PCIF1 knockout HeLa cells, or wild type parental cells, were infected with VSV at a MOI of 3. Viral RNA was radiolabeled and analyzed as in Fig 1 (representative images; n=3). **(B):** Add-back of PCIF1, but not a catalytically inactive mutant PCIF1_SPPG_ restores m^6^A_m_ on VSV mRNA. HeLa PCIF1 KO cells stably expressing 3X-FLAG-PCIF1, or 3X-FLAG-PCIF1_SPPG_ were infected with VSV and RNA radiolabeled, digested, and 2D-TLC performed as in A (representative images; n=3). **(C):** PCIF1 N6-methylates VSV mRNA *in vitro*. Purified VSV mRNA, transcribed *in vitro* from viral particles in the presence of [^32^P]-α-ATP, was used as template for *in vitro* methylation with 50 nM purified PCIF1, or PCIF1_SPPG_. Following hydrolysis the products were visualized by 2D-TLC and phosphorimaging as in Fig 1 (representative images; n=3).

### N7-guanosine methylation is dispensable for PCIF1 modification of VSV mRNA

Substrates for modification by PCIF1 require the presence of a methylated m^7^G mRNA cap structure (11, 12), and capped RNA lacking m^7^G serve as poor substrates for PCIF1 *in vitro* (14). The obligatory sequential methylation of cellular mRNAs at m^7^G and subsequent ribose-2’-O positions (1) precludes tests of the importance of the 2’-O methylation alone in PCIF1 modification of mRNA. By contrast the mRNA cap methylation reactions of VSV, and by inference other NNS RNA viruses, proceed in the opposite order. Experiments conducted with viral mutants and by reconstitution of cap-methylation *in vitro* demonstrate that cap ribose-2’-O methylation precedes and facilitates the subsequent guanine-N-7 methylation (28), allowing us to explore the impact of 2’-O methylation alone on PCIF1 modification of VSV mRNAs. Messenger RNA synthesized in cells by the viral mutant VSV-L_G1670A_ contains primarily GpppA_m_ mRNA caps (29), which serve as effective substrates for PCIF1 (Fig 3A). In agreement with analysis of viral mRNA made in cells, mRNA transcribed from purified VSV-L_G1670A_ virions are fully methylated by PCIF1 *in vitro* (Fig 3B, S3). Messenger RNA synthesized by a second viral mutant, VSV-L_G4A_, that produces unmethylated GpppA cap structures (29), was not modified by PCIF1 (Fig 3C). A low level of m^6^A observed following hydrolysis of RNA produced by VSV-L_G4A_ is consistent with modification at internal positions of the mRNA (Fig 1B). In agreement with this finding, viral mRNA synthesized by VSV in the presence of the methylation inhibitor *S*-adenosyl-homocysteine (SAH), is poorly methylated by PCIF1 (Fig 3D, S3). Taken together, this set of results demonstrate in contrast to cellular mRNA modification, VSV mRNA requires ribose-2’-O methylation and not guanine-N-7 methylation.

**Figure 3:**
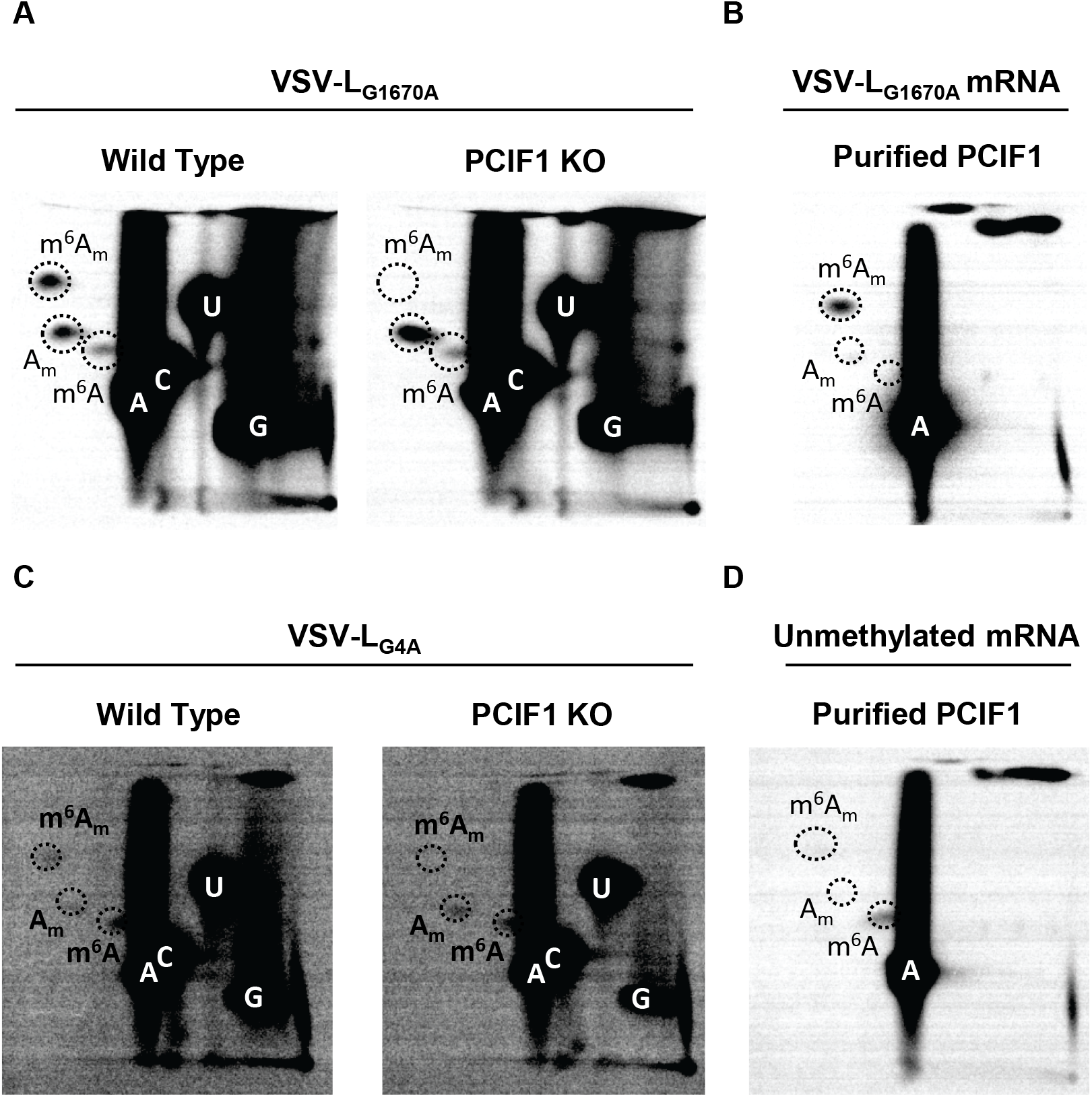
Effect of mRNA cap methylation on PCIF1 modification of VSV mRNA. **(A):** The indicated 293T cells were infected with VSV-L_G1670A_ at a MOI of 3 and viral RNA was radiolabeled, extracted and analyzed by 2D TLC as in Fig 2A (representative images, n=3). **(B):** Messenger RNA synthesized *in vitro* by VSV-L_G1670A_ mRNA was incubated with purified PCIF1 and analyzed as in Fig 2C (representative image, n=3). **(C):** The indicated 293T cells were infected with VSV-L_G4A_ as in panel A (representative images, n=3). **(D):** Messenger RNA synthesized by VSV *in vitro* in the presence of 200 μM SAH, was used as substrate for PCIF1 *in vitro* and analyzed as in panel B (representative image, n=3)

### Effect of m^6^A_m_ on viral growth, mRNA stability and translation

To determine whether PCIF1 modification influences VSV mRNA stability, we compared the decay of modified and unmodified transcripts. Briefly, mRNA was specifically isolated from infected PCIF1 KO cells, *in vitro* methylated with purified PCIF1 where indicated, transfected into uninfected cells, and isolated at the indicated time points. Analysis of the isolated mRNA by electrophoresis on acid-agarose gels demonstrates that viral mRNA stability is unaffected by the presence of m^6^A_m_ (Fig 4A, two-way ANOVA p>0.4). The stability of each individual mRNA appeared unaffected by PCIF1 dependent modification to m^6^A_m_ when transfected into either wild type or PCIF1 knockout cells (Fig S4).

**Figure 4:**
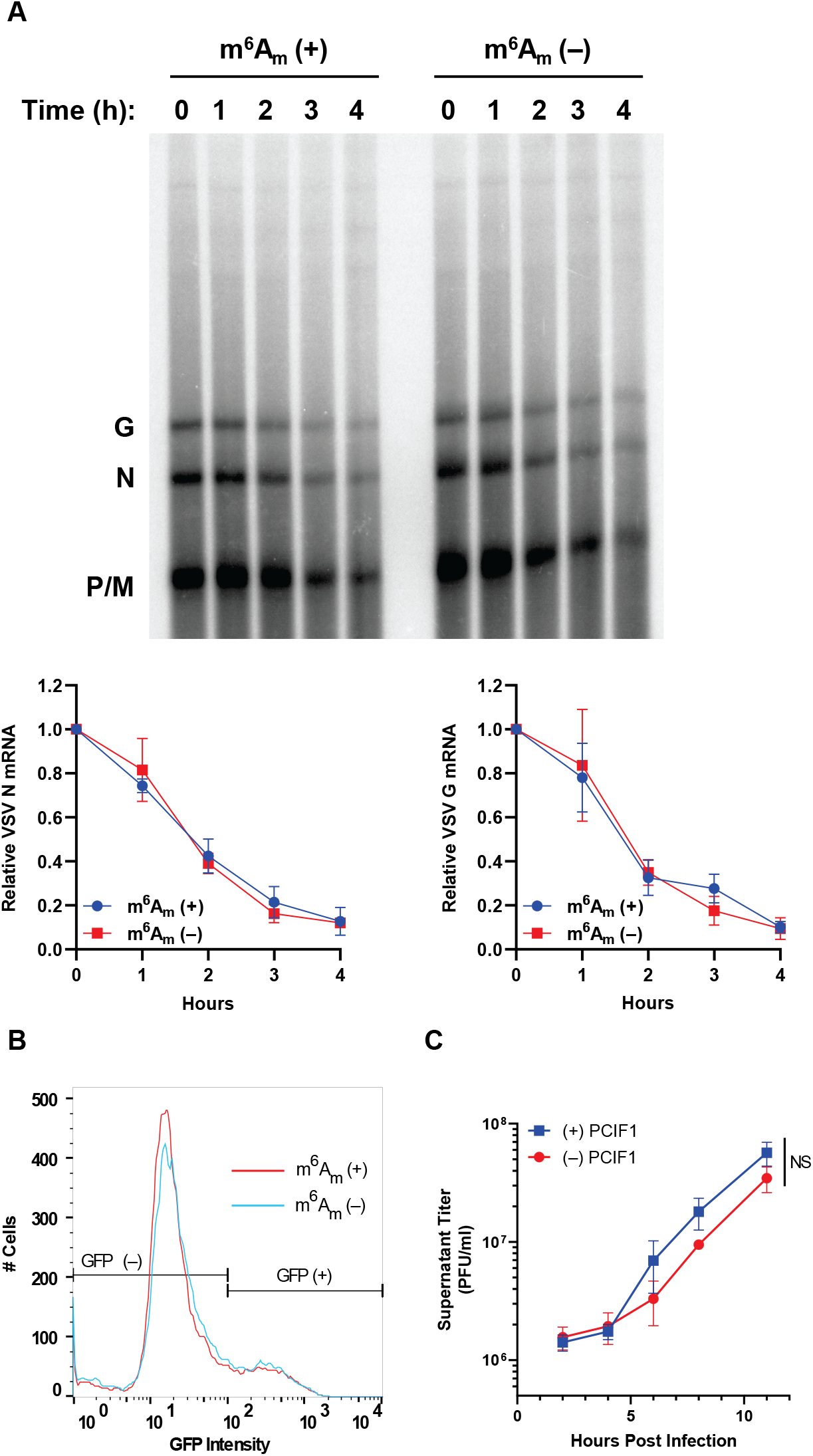
Effect on m^6^A_m_ on viral mRNA translation and stability. **(A-B)** VSV mRNA isolated from PCIF1 KO 293T cells and methylated *in vitro* with PCIF1 prior to transfection of 500 ng of RNA into PCIF1 KO HeLa cells and assessed for **(A):** mRNA stability by extraction from cells at the indicated time post-transfection and analyzed by electrophoresis on acid-agarose gels. A representative image is shown along with quantitative analysis of the abundance of the N and G mRNAs (n=3, +/−SD, 2-way ANOVA p>0.4). **(B)** mRNA translation by measurement of GFP positive cells and their intensity by flow cytometry (n=3, 0.95>p>0.18, student’s t-test). **(C)** Viral replication assessed in PCIF1 KO HeLa cells expressing 3X-FLAG-PCIF1 or an empty vector infected at a MOI of 3. Viral titers were determined by plaque assay on Vero cells at the indicated time post inoculation. (n=3, +/−SD, NS – p>0.08, student’s t-test, statistics shown are for the 11 h timepoint).

To measure whether mRNA translation was affected by the presence of m^6^A_m_, we measured the expression of a viral encoded eGFP reporter gene following transfection of methylated or unmethylated mRNA into PCIF1 KO cells. Measurement of eGFP by flow cytometry at 7 h post transfection reveals that neither the fraction of positive cells nor the fluorescence intensity was significantly altered by the presence of m^6^A_m_ (Fig 4B, S5A). Similar findings on stability and translation were obtained using a luciferase reporter (Fig S5B-C). To further verify that m^6^A_m_ does not impact the translation of VSV mRNA we co-transfected mRNA from two VSV reporter viruses encoding firefly (Luc) or renilla (Ren) luciferase with opposing methylations. In this competitive translation experiment the ratio of firefly and renilla translated in transfected cells was unaffected by the presence of m^6^A_m_ irrespective of which reporter virus mRNA was modified (Fig S5D-E). Collectively, these results demonstrate that translation of VSV mRNA is unaffected by the presence of m^6^A_m_.

As neither viral mRNA translation nor stability were altered by the presence of m^6^A_m_, we next compared the kinetics of viral growth in PCIF1 knockout or addback cells. Cells were infected at a MOI of 3 and viral titers determined by plaque assay at various times post infection. The kinetics of viral growth were also unaffected in cells lacking PCIF1 (Fig 4C, S6). Collectively these data demonstrate that despite extensive modification of the viral mRNAs by PCIF1, viral replication and gene expression are unaltered by this modification.

### Interferon-β treatment uncovers a role for PCIF1 in the host response to infection

As cap-methylation at the 2’-O position of the first nucleotide helps distinguish self from viral RNA during infection (4) we examined whether PCIF1 modification of mRNA plays a similar role. To examine whether cap-proximal m^6^A_m_ helps counter host cell antiviral responses, we measured how PCIF1 affects the IFN-β mediated inhibition of virus growth. Treatment of cells with IFN-β prior to infection uncovered a PCIF1 dependent attenuation of the antiviral effect (Fig 5A, S6). Infection of cells by a VSV-reporter virus that expresses firefly luciferase, confirmed that PCIF1 attenuates the suppressive effect of IFN-β on viral gene expression at the RNA and protein levels in a single-round of infection (Fig 5B, S7). This result suggests that the effects of PCIF1 are restricted to steps of the viral replication cycle up to and including gene expression. To eliminate viral entry as a possible contributor, we transfected ribonucleoprotein cores purified from VSV-Luc into cells, thereby bypassing viral entry. Pretreatment of cells with IFN-β was still accompanied by augmented inhibition of gene expression in cells lacking PCIF1 (Fig 5C). This result demonstrates that the IFN-β mediated suppression of VSV gene-expression is enhanced in cells lacking PCIF1.

**Figure 5:**
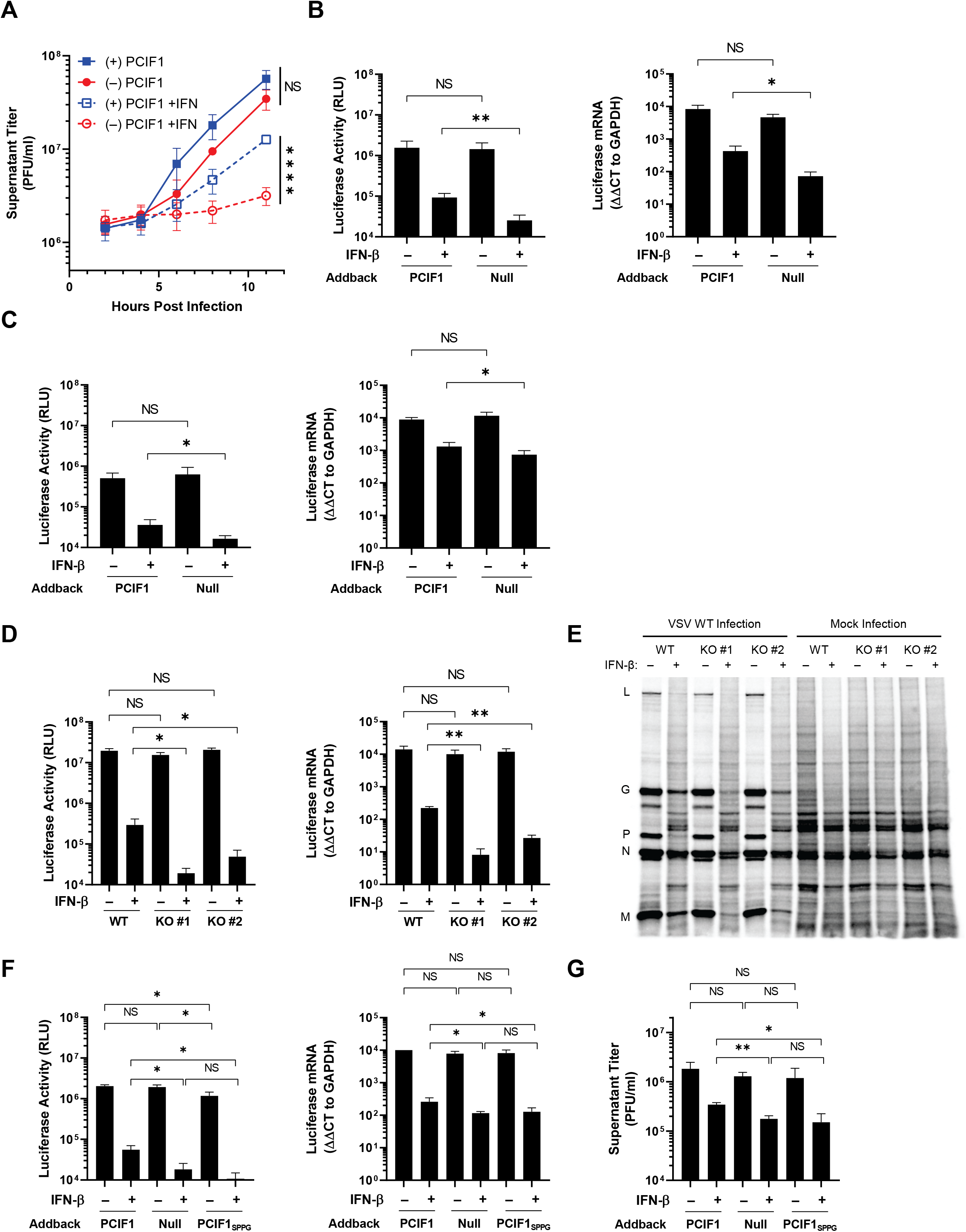
Effect of IFN-β pretreatment of cells on viral infection. **(A-C):** PCIF1 KO HeLa cells reconstituted with PCIF1 or an empty vector were pretreated with vehicle (0.1% BSA) or 500 U ml^−1^ of interferon-β for 5h and infected with the indicated VSV at a MOI of 3. **(A):** VSV viral titer was determined at the indicated timepoints by plaque assay on Vero cells (n=3, +/− SD, NS – p>0.08, **** p<0.0001, student’s t-test). **(B):** Cells were infected with a VSV-Luciferase reporter and luciferase activity measured by luminometer at 6 hpi (n=4, +/−SD, NS – p>0.80, ** – p<0.01, student’s t-test). Quantitative RT-PCR analysis of luciferase mRNA (normalized to GAPDH, n=4, +/−SD, NS – p>0.05, * – p<0.05, student’s t-test). **(C):** As in B, except cells were transfected with 500 ng ribonucleoprotein cores of VSV-Luc. (n=4, +/−SD, NS – p>0.19, * – p<0.05, student’s t-test). **(D):** The indicated A549 cells were pretreated with vehicle (0.1% BSA) or 500 U ml^−1^ IFN-β for 5 hours prior to infection with VSV-Luc at a MOI of 5. Cells were lysed 6 hpi and luciferase activity determined as in panel B (n=4, NS – 0.50>p>0.05, * – p<0.05, student’s t-test) and mRNA levels verified by qRT-PCR as in panel B (n=3, NS – 0.47>p>0.23, * – p<0.05, ** - p<0.01, student’s t-test). **(E):** As in panel D, with infection by VSV assessed by metabolic incorporation of [^35^S]-met and [^35^S]-cys into viral proteins as described in methods. Proteins were analyzed by SDS-PAGE and visualized by phosphorimager (representative image, n=3). **(F):** As in B, cells were reconstituted with PCIF1, PCIF1_SPPG_ or empty vector (n=3, +/− SD, NS – 0.83>p>0.05, * – p<0.05, student’s t-test). **(G)**: As in F, except cells were infected with VSV and viral titers measured at 11 hpi by plaque assay (n=3, +/− SD, NS – 0.83>p>0.29, * – p<0.05, student’s t-test).

The antiviral response in HeLa cells is partly attenuated (30), therefore we examined whether loss of PCIF1 results in a similar IFN-β dependent inhibition of viral replication in A549 cells. We confirmed that VSV mRNAs were also N6-methylated by PCIF1 in these cells (Fig S8). Pretreatment of A549 cells with IFN-β revealed that, in the absence of PCIF1, viral gene expression was suppressed an additional 10-fold as evident by levels of viral mRNA and protein (Fig 5D). Measurements of specific viral proteins following metabolic incorporation of [^35^S]-met and [^35^S]-cys followed by analysis of proteins on SDS-PAGE demonstrates that the 3 most abundant viral proteins, N, M and G are further suppressed in cells lacking PCIF1 (Fig 5E, S9), but cellular translation in uninfected cells is unaffected (Fig 5E, S9).

To rule out the possibility of N6-methylation independent activities of PCIF1 mediating this effect, we examined infection in cells expressing a catalytically-inactive mutant of PCIF1, PCIF1_SPPG_ (Fig 2B). Both PCIF1_SPPG_ addback and PCIF1 knockout equivalently augmented the effect of IFN-β on VSV-luciferase gene expression (Fig 5F) and on viral growth (Fig 5G). Collectively, the above experiments reveal that loss of PCIF1 or its ability to synthesize m^6^A_m_ augments the suppressive effect of IFN-β on VSV gene expression, suggesting that m^6^A_m_ methylation of viral mRNAs protects against the otherwise antiviral effects of the IFN-mediated innate immune response.

## Discussion

The major finding of this study is the identification of a role for PCIF1-mediated m^6^A_m_ methylation in the type I interferon response to VSV infection. We demonstrate that loss of PCIF1 enhances the sensitivity of viral replication to pretreatment of cells with IFN-β by affecting VSV gene expression. PCIF1 is necessary and sufficient for modification of VSV mRNA to yield cap-proximal m^6^A_m_, and in contrast to cellular mRNA modification, viral mRNAs require prior ribose-2’-O but not guanine-N-7 methylation of the cap-structure. The most parsimonious explanation of our results is that the PCIF1 dependent modification of viral mRNA cap-structures to m^6^A_m_ serves to dampen an IFN-β mediated suppression of gene expression. Mechanistically, how this occurs was not resolved by the present study but we posit that this requires discrimination of modified from unmodified RNA by an interferon stimulated gene (ISG).

Precedent for a role of mRNA cap modifications in the antiviral response already exists. The RIG-I dependent recognition of a 5’ triphosphate is suppressed by the presence of an mRNA cap-structure (4, 6), and ribose 2’-O methylation of the cap-structure inhibits the ability of an ISG, IFIT1, to suppress translation of mRNA (4, 7). The PCIF1 dependent modification of VSV mRNA cap-structures may work by a similar mechanism by helping viral mRNA appear more host like. Additional work will be necessary to define whether an ISG is required to discriminate between m^6^A_m_ modified and unmodified cap-structures. We suggest that it is unlikely that IFIT1 functions in this discrimination based on its known recognition of m^7^GpppA cap-structures (31) and the requirement for 2’-O modification of VSV mRNA for their subsequent PCIF1 modification (32).

Although we establish a role of PCIF1 and m^6^A_m_ in the IFN-β mediated suppression of VSV gene expression, and demonstrate that viral mRNAs are modified by PCIF1, modification of cellular mRNA may also play a role. Cap proximal m^6^A_m_ inhibits the host mRNA decapping enzyme DCP2 (17), which may alter stability of cellular mRNAs including those induced on treatment of cells with IFN-β (33). This seems unlikely to account for the effects we observe on VSV infection, as the DCP2 dependent decapping and degradation of cellular mRNA would be expected to increase in cells lacking PCIF1, likely dampening rather than augmenting the effect of IFN-β treatment. If the antiviral response is due to m^6^A_m_ modification of cellular mRNA this contrasts with the consequences of 2’-O methylation of the mRNA cap structure of ISG mRNAs which enhances their expression (34). We are also mindful of the possibility that PCIF1 may have unknown functions in the cell beyond N6-methylation of mRNA. Insects, including *Drosophila*, express an ortholog of PCIF1 that associates with the phosphorylated CTD of PolII, but is catalytically inactive as an RNA N6-methyltransferase (35). As the catalytic activity of PCIF1 is required for attenuation of the antiviral response we also find this explanation unlikely.

The substrate requirements for PCIF1 modification of VSV mRNA differ to those previously shown in cellular mRNA (11, 14). Specifically, we found that guanine-N7-methylation was dispensable for N6-methylation, and that ribose 2’-O methylation was required. Although we do not understand why this is the case, we recapitulate the substrate specificity *in vitro* making it unlikely that this distinction reflects the cytoplasmic modification of viral mRNA rather than the nuclear modification of cellular mRNA. This altered specificity for modification of the VSV mRNA coupled with the altered recognition specificity of the VSV cap methylation machinery - which requires 2’-O methylation prior to guanine-N-7 methylation – raise the possibility that the structure of the 5’ end of VSV mRNAs leads to the altered specificity (29).

The finding that PCIF1 and m^6^A_m_ affect the antiviral response raises the question of why modify cap-proximal A and not other cap-proximal bases. More cellular mRNAs initiate with G than A (16), but guanosine is typically only methylated at the 2’O position in mRNA (36). This is likely because O6-methylation of guanosine (m^6^G) has been shown to have a large fitness cost. Its presence in DNA is highly mutagenic though pairing with thymidine during DNA replication, and when present in RNA, it causes incorrect ribosome decoding (37) and a 1000-fold decrease in the peptide-bond formation rate (38). N6-methylation in adenosine, by contrast, has minimal effect on these processes (37, 38).

The importance of m^6^A_m_ for other viruses has not been examined. As expected, the mRNAs of DNA viruses which rely on host RNA polymerases for transcription, including adenoviruses (39, 40), simian virus 40 (41), herpes simplex virus 1 (42), and polyomaviruses (43) contain m^6^A_m_ (1, 3). Vaccinia virus, which replicates in the cytoplasm, also produces m^6^A_m_ containing mRNA (1, 3, 44), likely through modification by PCIF1. It remains largely unexplored for other RNA viruses which produce mRNAs that initiate with A. The evolution by many viruses of their own capping machinery also begs the question of whether viruses have evolved a PCIF1-like cap modifying enzyme, particularly given the potential advantage in the face of an antiviral response. Additional studies with VSV and other viruses will be required to fully define the role of PCIF1 in the host response to infection. Rabies virus, for example, also produces 5 mRNAs that initiate with a similar sequence to those of VSV and therefore are likely to be modified by PCIF1. VSV and rabies antagonize the innate immune response through distinct mechanisms suggesting that this comparison may help further illuminate the role of PCIF1 in the host response to infection (45–47).

## Materials and methods

### Cells

HEK293T, HeLa, A549, Vero CCL81, and BsrT7/5 cells were maintained in humidified incubators at 37 ºC and 5% CO_2_ in DMEM supplemented with L-glutamine, sodium pyruvate, glucose, (Corning #10013CV) and 10% fetal bovine serum (Tissue Culture Biologicals #101). Generation of HEK293T and HeLa *PCIF1-*knockout cell lines and addbacks were previously described (12), and A549 *PCIF1*-knockout cell lines were generated and verified using these same methods. Cells were tested regularly using the e-Myco PLUS PCR kit (Bulldog Bio #2523348).

### Viruses

VSV (as rescued from an infectious cDNA clone of VSV, pVSV(1)+), VSV-L_G1670A_, VSV-L_G4A_, VSV-luciferase, VSV-RenillaP, and VSV-eGFP have been described previously (29, 48–51). Viruses were propagated in BsrT7/5 cells.

### Radiolabeling of mRNA

Cap-proximal nucleotides were radiolabeled as previously described (17). For specific radiolabeling of VSV mRNA, cells were infected or mock infected with VSV at a MOI of 3 in serum/phosphate-free DMEM (Gibco #11971-025). 10 μg ml^−1^ Actinomycin D (Sigma #A5156) was added to halt cellular transcription at 2.5 hpi, and at 3 hours post infection, 100 μCi ml^−1^ [^32^P] phosphoric acid added to label newly synthesized RNA (Perkin Elmer #NEX053H). RNA was harvested in Trizol (Thermofisher #15596018) and poly(A)+ mRNA selected using the NEB Magnetic mRNA Isolation Kit (NEB S1550S).

### Identification of methylated nucleotide levels by two-dimensional thin layer chromatography (2D-TLC)

2 μg mRNA suspended in 6 μl RNAse-free H_2_O was digested with 2 units of nuclease P1 (Sigma N8630) for 3 h at 37 ºC. The volume was then increased to 20 μl and RNA further digested with 2 units of Cap-Clip Acid Pyrophosphatase for 3 h (Cell Script #C-CC15011H) in the manufacturer’s buffer. 2D-TLC was performed as previously described (27). Plates were developed in the first dimension with 5 parts isobutyric acid (Sigma #I1754) to 3 parts 0.5 M ammonia (VWR #BDH153312K) for 14 h, and in the second dimension with a solvent of 70 parts isopropanol, 15 parts hydrochloric acid, and 15 parts water for 20 h. RNA species were positively identified by UV-shadowing (254 nm) of co-spotted (non-radioactive) commercially available standards (5’ monophosphate forms). The standards A_m_ and m^6^A_m_ 5’ monophosphate were generated by digesting their triphosphate forms (TriLink N-1015 and N-1112) with 1 unit Apyrase (NEB M0398).

### In vitro transcription of VSV mRNA and methylation with PCIF1

VSV or VSV-L_G1670A_ mRNA was synthesized *in vitro* as previously described with 30 μCi [^32^P]-α-ATP per reaction (29, 52). RNA was extracted in trizol, poly(A) selected, and extracted in trizol again to concentrate the samples. Purified recombinant GST-PCIF1 was generated and used to in vitro methylate 225 ng of this mRNA as previously described (12).

### Purification, selection, and in vitro methylation of VSV mRNA from cells

HEK 293T *PCIF1* KO cells were infected at a MOI of 10 for 7 h. RNA was extracted in trizol, and VSV mRNAs selected using a biotinylated oligo against the conserved stop and poly(A) sequence present at the 3’ end of VSV mRNAs (“Biotin-VSVstop”). 1.5 nmol oligo was annealed to this mRNA by incubating at 65 ºC for 5 min, followed by cooling on ice for 5 min, and complexes isolated by pulldown with NEB Streptavidin Magnetic Beads (NEB #S1420, manufacturer’s protocol). Following cleanup by trizol extraction, mRNA was purified further using the NEB poly(A) magnetic kit as above, and trizol extracted again. This stock of RNA from PCIF1 KO cells was then *in vitro* methylated (or mock methylated) with purified PCIF1 as above.

### Determination of VSV mRNA stability

Biotin-VSVstop selected mRNAs were transfected into HeLa WT or *PCIF1* KO cells. RNA was transfected into separate wells of a 24 well plate (2×10^5^ cells; 500 ng RNA per well) using Lipofectamine 2000 (Thermofisher #11668019), media was changed at 3 h post transfection, and wells harvested in trizol at 1 h intervals for 4 h. Extracted mRNAs were separated by electrophoresis on acid-agarose gels, which were dried and exposed to a phosphor screen. VSV mRNAs levels were quantified using ImageQuant version 8.2, and normalized to the 0 h timepoint.

### Transfection and flow-cytometry analysis of translation of VSV mRNA

500 ng Biotin-VSVstop selected VSV mRNAs were transfected into 2×10^5^ HeLa WT or *PCIF1* KO cells using Lipofectamine 2000. At 6 hours post transfection, cells were trypsinized, and washed and resuspended in PBS. Half the cells were analyzed for GFP expression by flow cytometry (BD FACS Calibur); GFP positive cells and mean fluorescence intensity of GFP were calculated in FlowJo (20,000 cells analyzed per replicate)

### Determination of VSV mRNA gene expression by luciferase luminescence and RT-qPCR

2×10^5^ cells (24 well format; transfection experiments) or 4×10^5^ cells (12 well format; IFN experiments) were lysed in 120 μl passive lysis buffer (Promega #E1941). Half the lysate was used to quantify luciferase protein (Promega Luciferase Assay System #E1501) using a Spectramax L luminometer with reagent injectors in technical triplicate. RNA was extracted from the other half of cells in trizol, and 1 μg reverse transcribed using SuperScript III (Invitrogen #18080044), oligo-dT primers (IDT #51011501), and RNase inhibitors (Promega #N2515). Real-time qPCR was performed using Fast SYBR Green (Thermofisher #4385612) in technical duplicate. Relative RNA was calculated as ∆∆CT (normalized to GAPDH) times 10^4^.

### Detection of Radiolabeled Samples

Gels were fixed in 30% methanol, 10% acetic acid, washed twice in methanol, and dried using a vacuum pump gel dryer. TLC plates were air dried. Dried gels or plates were then exposed to a phosphor screen and scanned on a Typhoon scanner.

### Growth Curve with IFN pretreatment

HeLa cells were pretreated with 500 U ml^−1^ IFN-β (Tonbo Biosciences 21-8699) or vehicle (0.1% BSA) for 5 h in serum free DMEM. Cells were washed, and infected with VSV at a MOI of 3 for 1 h in serum free DMEM. After 1 hour, the inoculum was removed, cells washed, and supplemented with 2% FBS. At 2, 4, 6, 8, and/or 11 hours post infection, 1% of the supernatant was removed and frozen at −80 ºC. After all samples had been collected, viral titers were determined by plaque assay on vero cells.

### Gene expression with IFN pretreatment

HeLa or A549 cells were pretreated with 500 U ml^−1^ IFN-β as above for 5 h, infected with VSV-Luc at a MOI of 3, and at 6 hours post infection, cells were lysed and processed for luciferase protein and mRNA quantitation. Alternatively, VSV-Luc RNPs were purified as previously described, and 50 ng transfected into HeLa cells instead of infection with virus.

### Metabolic Radiolabeling of Protein

4×10^5^ A549 cells were pretreated with 500 U ml^−1^ IFN-β as above for 5 h, and infected or mock infected with wild type VSV at MOI 5. At 5 hours post infection, cells were washed, and the media changed to methionine/cysteine free DMEM (Gibco #21013-024). After 40 minutes of starvation, cells were pulse labeled with 30 μCi ml^−1^ [^35^S] methionine (Perkin Elmer #NEG009T) and 30 μCi ml^−1^ [^35^S] cysteine (Perkin Elmer #NEG022T) for 20 minutes. Cells were then lysed in SDS sample buffer and run on a low-bis 10% SDS-PAGE gel. Protein translation was determined by phosphorimaging as above. Equal loading was determined by staining with 0.25% Coomassie Brilliant Blue G-250.

### Data analysis and replicates

All experiments were performed with n=3 or n=4 biological replicates (as indicated). Each qPCR biological replicate is the average of technical duplicates from the same sample. All qPCR biological replicates (except Fig 5F) were run and analyzed on the same plate, enabling a standard deviation to be calculated for all samples. Each luciferase biological replicate is the average of technical triplicates from the same sample. Statistical tests were performed in Microsoft Excel, graphs were generated in Graphpad Prism 8.

## Acknowledgments

Thanks to members of the S.P.J.W. laboratory who provided insightful discussions and advice. This project was supported by NIH Grants F31AI138448 (to M.A.T), T32AI007245 (to S.P.J.W.), AI059371 (to S.P.J.W.), DP2AG055947 (to E.L.G), and R21HG010066 (to E.L.G).

---

**Figure S1:**
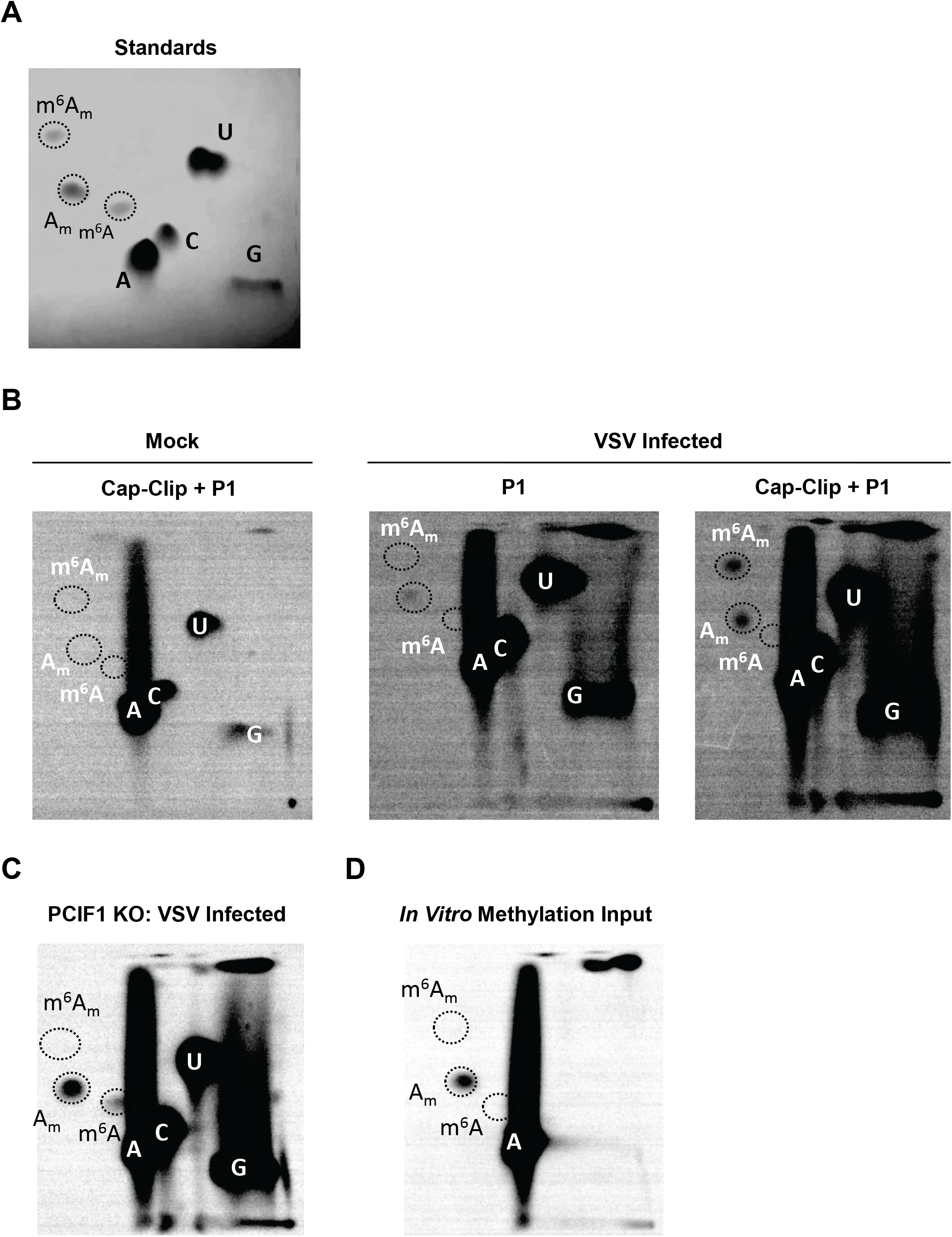
VSV mRNAs in HeLa cells contain a 5’ m^7^Gpppm^6^A_m_ cap-structure. **(A):** Chemical standards used to identify nucleotide species visualized by UV shadowing (254 nm). **(B):** HeLa cells were infected with VSV at a MOI of 3, cellular transcription halted by adding 10 μg ml^−1^ actinomycin D at 2.5 hpi, and viral RNA metabolically labeled with 100 μCi ml^−1^ [^32^P] phosphoric acid from 3-7 hpi. RNA was extracted, poly(A) selected and incubated with the indicated nucleases and the products resolved by 2D-TLC and detected by phosphorimaging. Wild type HeLa cells (representative image; n=3). **(C):** 293T PCIF1 KO cells **(D):** In vitro transcribed VSV mRNA (input to Fig 2C).

**Figure S2:**
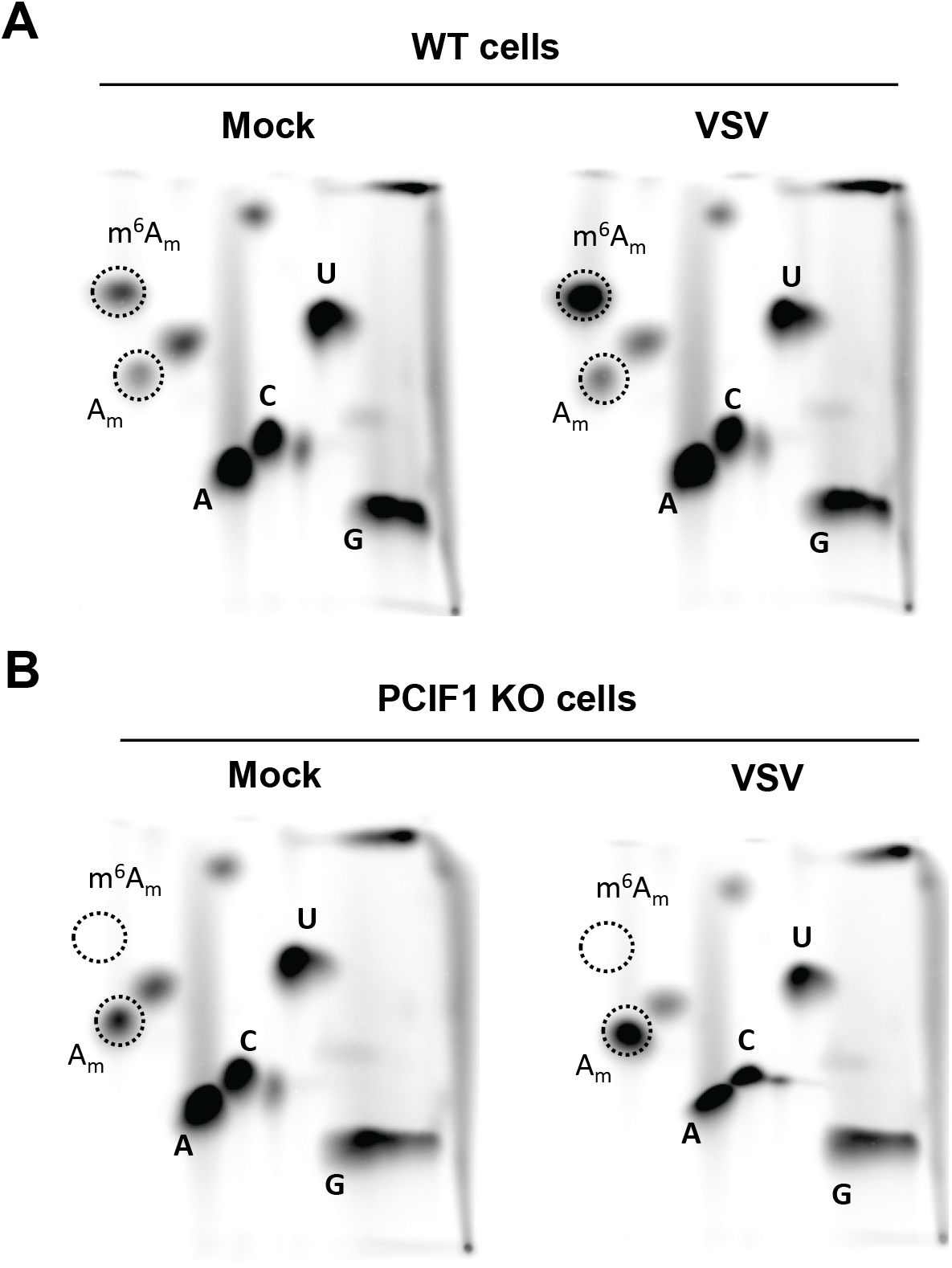
PCIF1 methylates viral mRNA. 293T cells were infected with VSV at a MOI of 3, poly(A)+ RNA purified at 6 hpi, and cap-proximal nucleotide identity determined by selective radiolabeling. Poly(A)+ RNA was decapped with Cap-Clip, and the exposed 5’ phosphate of the cap-proximal nucleotide was radiolabeled with [^32^P] γ-ATP by sequential treatment with shrimp alkaline phosphatase and polynucleotide kinase. Hydrolyzed nucleotide monophosphates were resolved by 2D-TLC and detected by phosphorimaging (representative images; n=3). **(A):** Parental wild type 293T cells. **(B):** PCIF1 knockout 293T cells.

**Figure S3:**
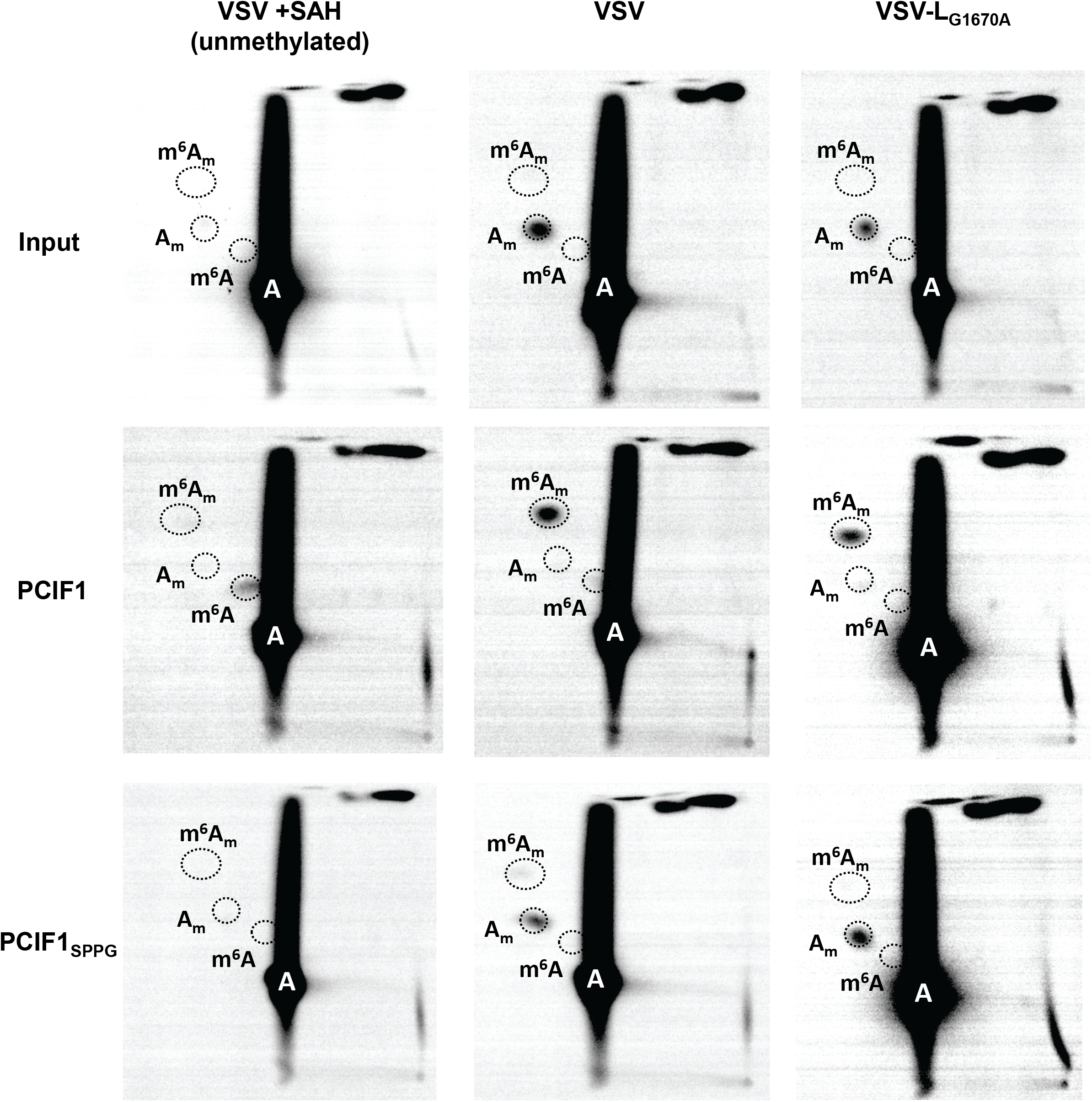
Cap-methylation requirements for i*n vitro* methylation of VSV mRNA. VSV mRNA was transcribed *in vitro* using purified virions from the indicated viruses used in Fig 3, and in the presence or absence of 200 μM SAH (a methylation inhibitor) as noted, followed by *in vitro* methylation with no enzyme (“input”), purified PCIF1, or purified PCIF1_SPPG_. 2D-TLC was performed on the products to determine the relative amounts of m^6^A_m_ and A_m_ present (representative images; n=3).

**Figure S4:**
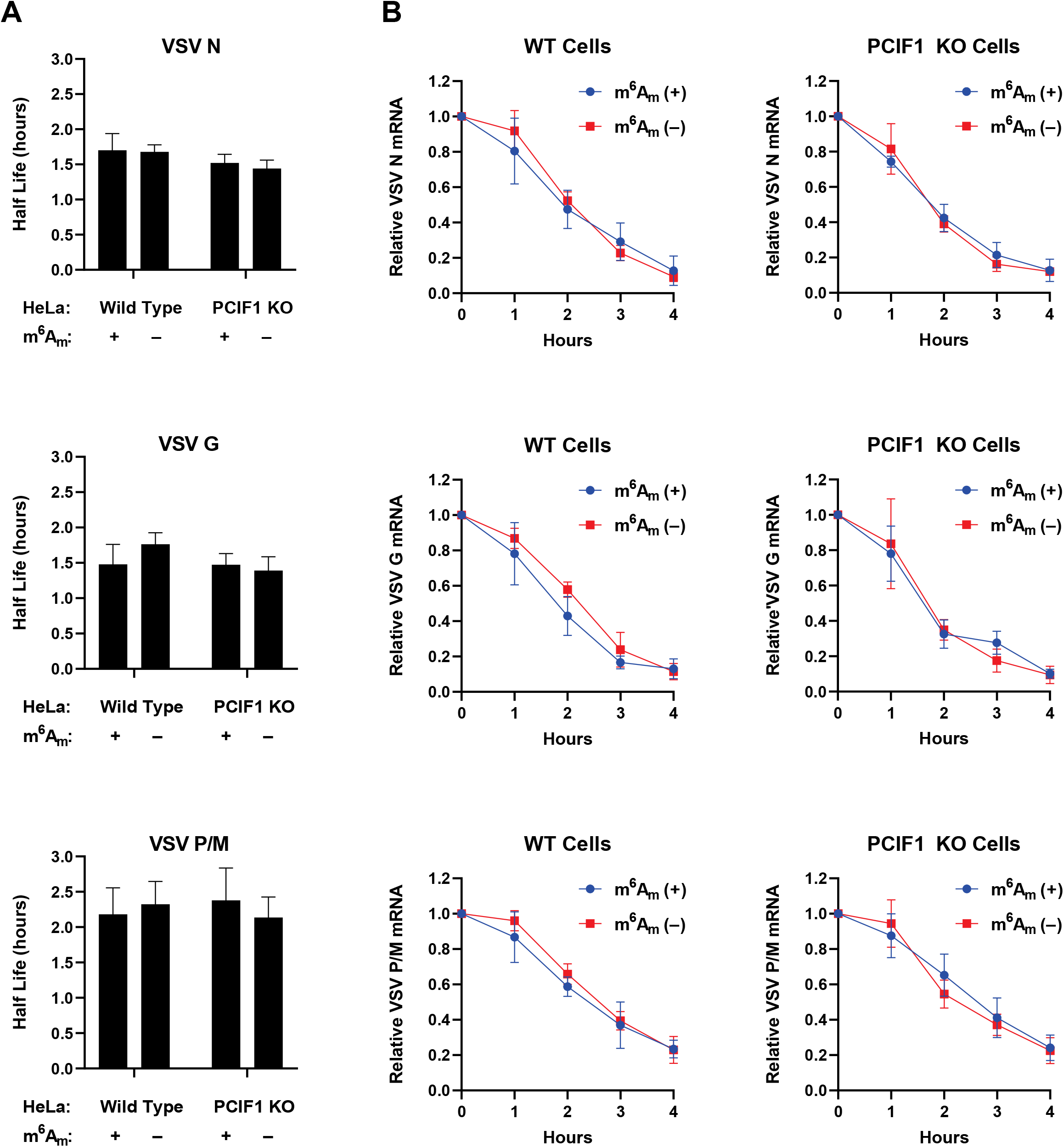
Effect of m^6^A_m_ on VSV mRNA stability. Purified stocks of VSV radiolabeled mRNA were generated by infecting 293T PCIF1 KO cells at a MOI of 10 with VSV as in Fig 1, followed by purification. Purified RNA (500 ng) was in vitro methylated with PCIF1, transfected into HeLa cells and RNA amounts assessed by re-extraction from cells at the indicated times followed by electrophoresis on acid-agarose gels and phosphorimaging (see Fig 4). **(A):** mRNA half life calculated from each decay curve (n=3, +/− SD). **(B):** Decay curves used to calculate half lives. There is no significant difference between any decay curve (2 way ANOVA, 0.75>p>0.4), or calculated half life (student’s t-test, 0.98 >p>0.06).

**Figure S5:**
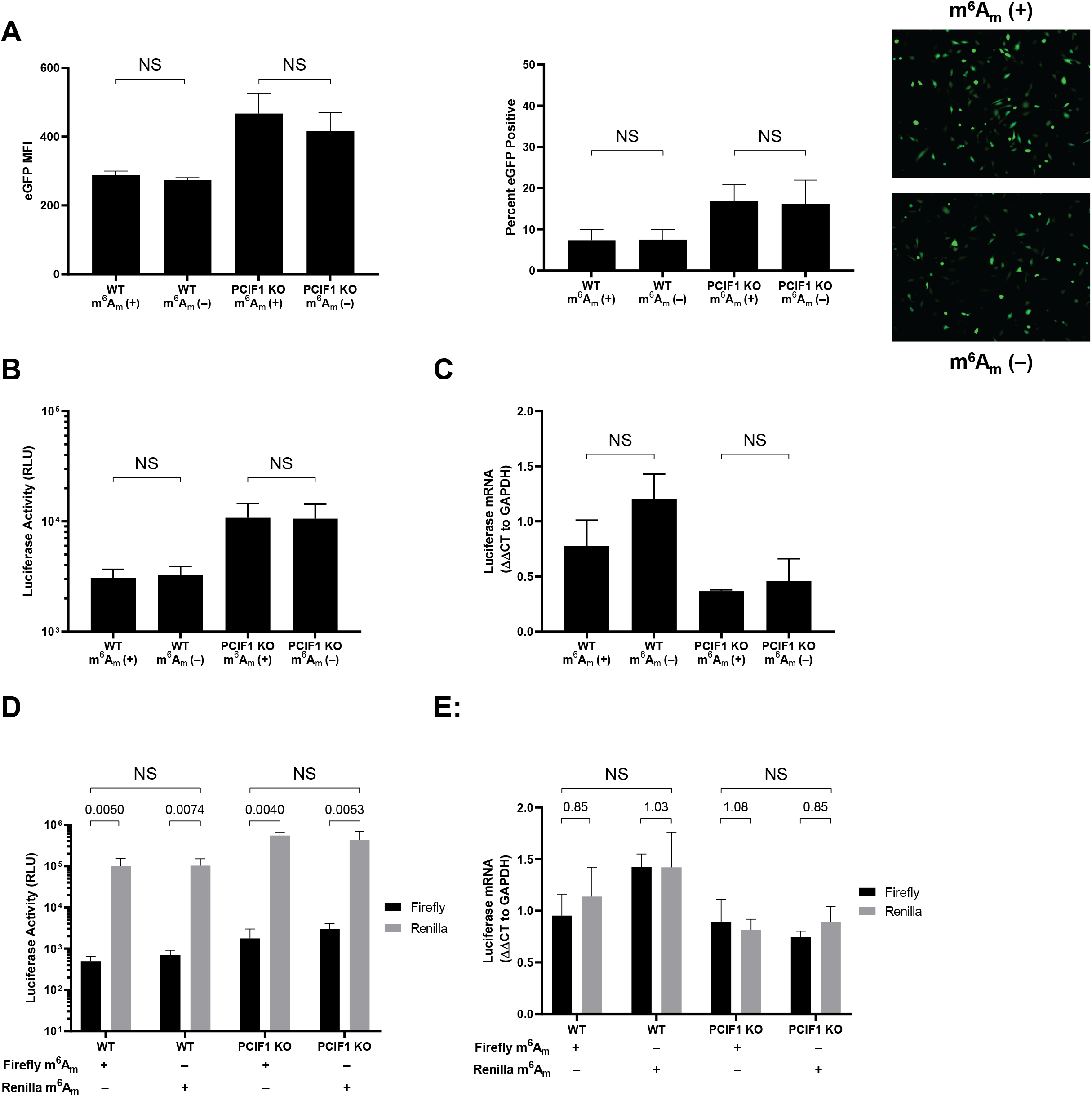
Effect of m^6^A_m_ on translation of VSV mRNA reporters. Purified stocks of VSV mRNA from the indicated virus were generated by infecting 293T PCIF1 KO cells at a MOI of 10 with VSV as in Fig 4, followed by purification of extracted RNA by poly(A) selection and a biotinylated oligonucleotide against the conserved VSV-stop sequence. 500 ng RNA was then mock or in vitro methylated with purified PCIF1, and transfected into HeLa wild type or PCIF1 knockout cells. **(A):** N6-methylation does not impact translation of a GFP reporter. VSV-eGFP mRNA was transfected into cells, and flow cytometry performed at 6 hpi. Mean fluorescence intensity (n=3, +/− SD, NS – p>0.18, student’s t-test,) and percent GFP positive cells (n=3, +/− SD, NS – p>0.89, student’s t-test) are shown, with a representative florescence microscopy image of transfected cells. **(B):** N6-methylation does not impact translation of a luciferase reporter. 500 ng VSV-luciferase mRNA with the indicated methylation was transfected into the indicated HeLa cells. Cells were lysed at 6 hpi, and luciferase levels were measured using a Promega Luciferase Assay kit (n=3, +/− SD, NS – p>0.70, student’s t-test). **(C):** RNA was extracted from lysate from (B), RT-PCR performed with oligo-dT primers, and qPCR performed for luciferase RNA (n=3, normalized to GAPDH, +/− SD, NS – p>0.08, student’s t-test). **(D):** Translation of an m^6^A_m_ (+) reporter does not outcompete a co-transfected m^6^A_m_ (–) reporter. 300 ng purified VSV-Luc (firefly) and VSV-RenP (renilla) mRNAs with opposing methylation status (m^6^A_m_ (+) firefly with m^6^A_m_ (–) renilla, and vice versa) were transfected into the indicated HeLa cells for 8 hours. Cells were lysed and luciferase levels of both reporters using a Promega Dual-Luciferase kit. Relative luminescence units (RLU) are shown (n=3, +/− SD, NS – 0.98 >p>0.11, student’s t-test). Ratios of Firefly to Renilla are shown above each condition. **(E):** No change in luciferase RNA levels from (D). RNA from (D) was extracted and qPCR performed as in C for Firefly and Renilla luciferase transcripts (n=3, normalized to GAPDH, +/− SD, 0.49>p >0.05, student’s t-test).

**Figure S6:**
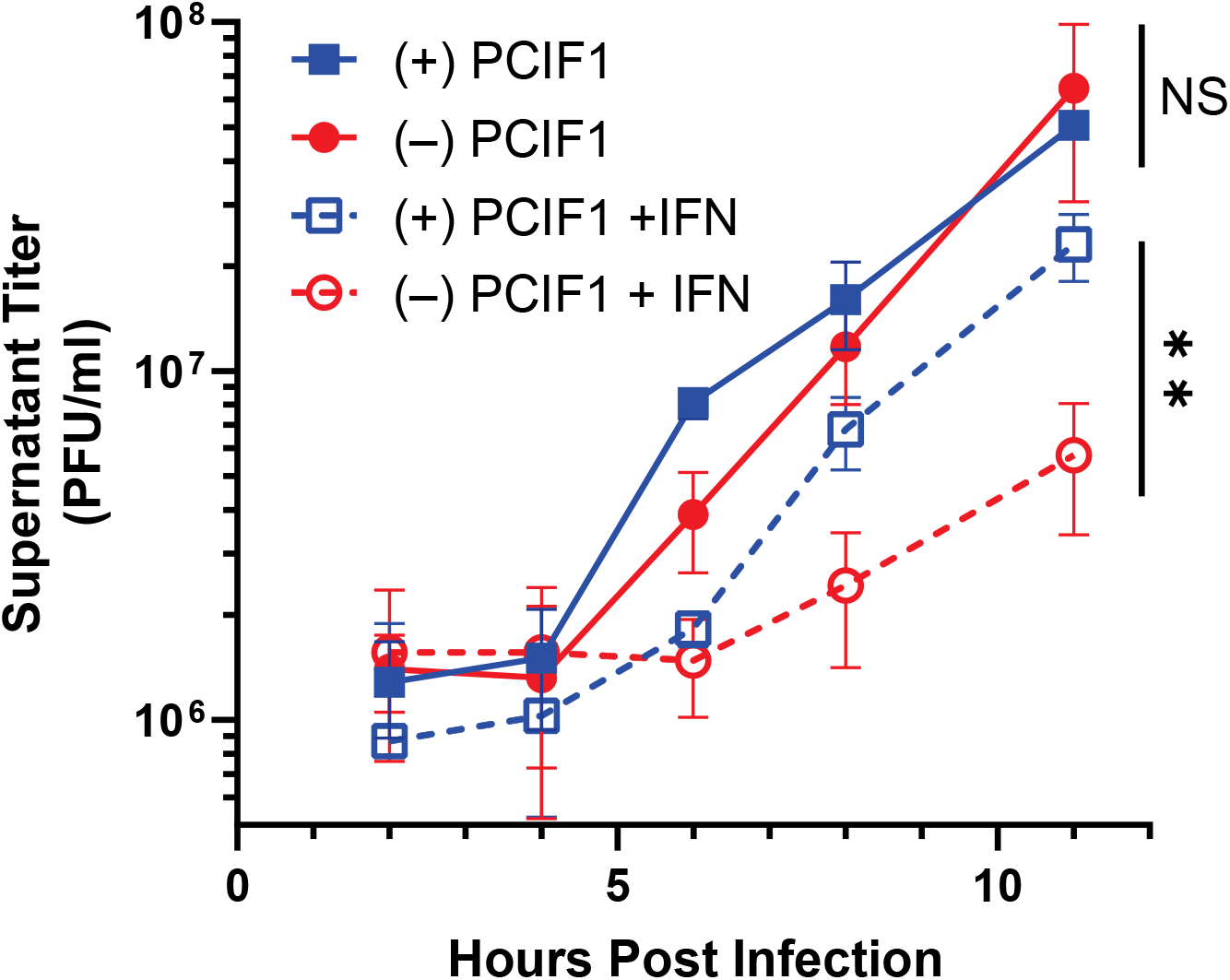
Effect of IFN-β pretreatment of cells on viral infection in a second *PCIF1*-addback clone. PCIF1 KO HeLa cells (different single cell clone from Fig 4C, 5A) reconstituted with PCIF1 or an empty vector were pretreated with vehicle (0.1% BSA) or 500 U ml^−1^ interferon-β for 5h. Treatment media was removed from the cells, followed by infection with VSV WT at a MOI of 3. After 1 hour, the inoculum was removed, cells washed, and initial treatment media added back to cells. At 2, 4, 6, 8, and 11 hpi, 1% of the supernatant was removed, and plaque assays performed on Vero cells to determine the titer of VSV in each sample. Growth curve of supernatant virus (n=3, +/−SD. NS – p>0.51, ** - p<0.01, student’s t-test. Statistics shown are for the 11h timepoint).

**Figure S7:**
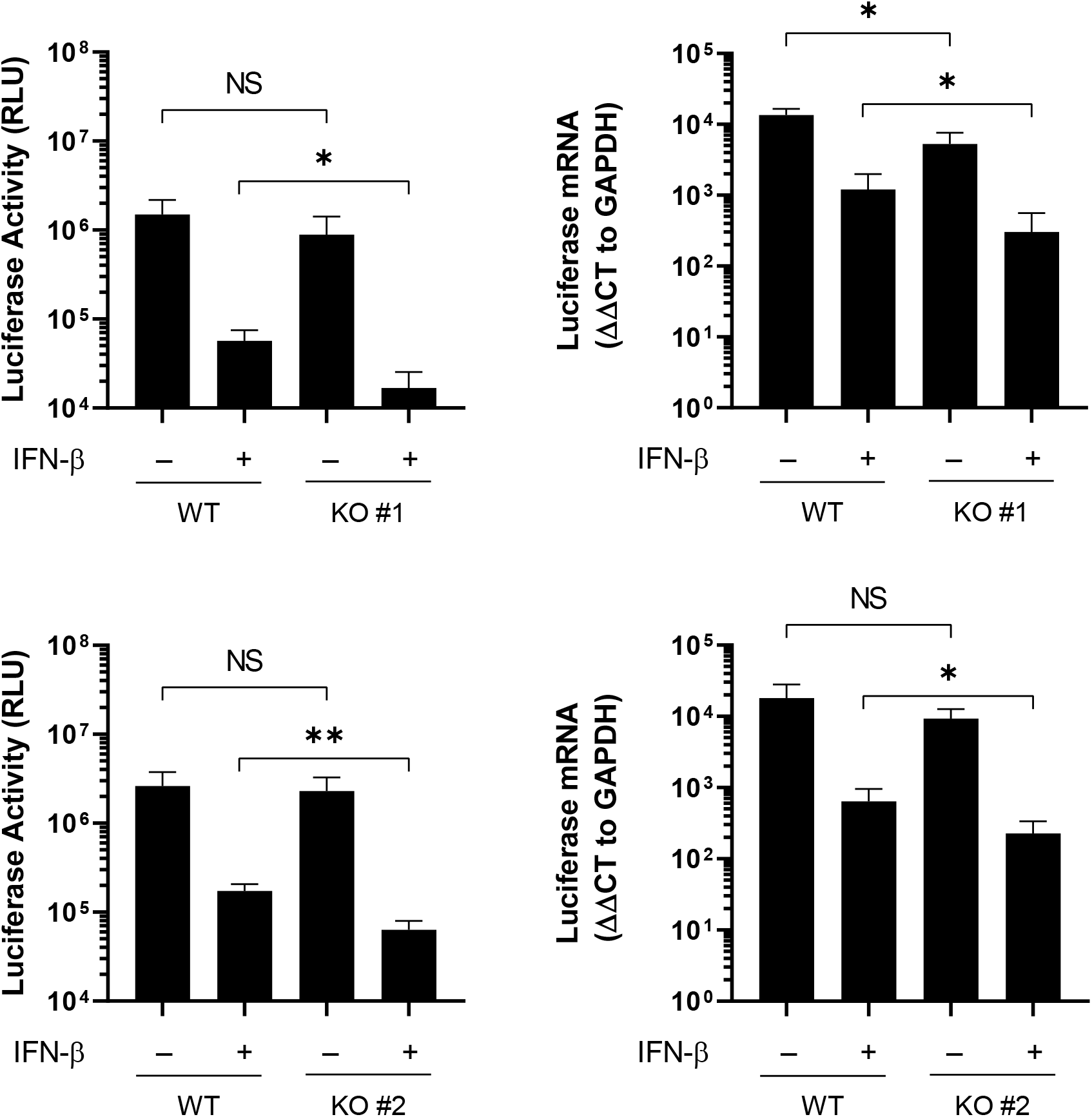
Effect of IFN-β pretreatment of cells on viral infection in multiple single-cell clones of PCIF1 KO cells. Wild type (parental) HeLa or PCIF1 KO cells (two independent clones) were pretreated with IFN-β for 5h, then infected with VSV expressing a luciferase reporter (VSV-Luc) at a MOI of 3. Cells were lysed at 6 hpi. Half the lysate was used to measure luciferase using a Promega Luciferase Assay kit (n=4, NS – p>0.20, * - p<0.05, ** - p<0.01, student’s t-test). RNA was extracted from the other half in Trizol and RT-PCR performed using oligo-dT, followed by qPCR for luciferase mRNA (normalized to GAPDH, n=4, NS – p>0.18, * - p<0.05, ** - p<0.01, student’s t-test).

**Figure S8:**
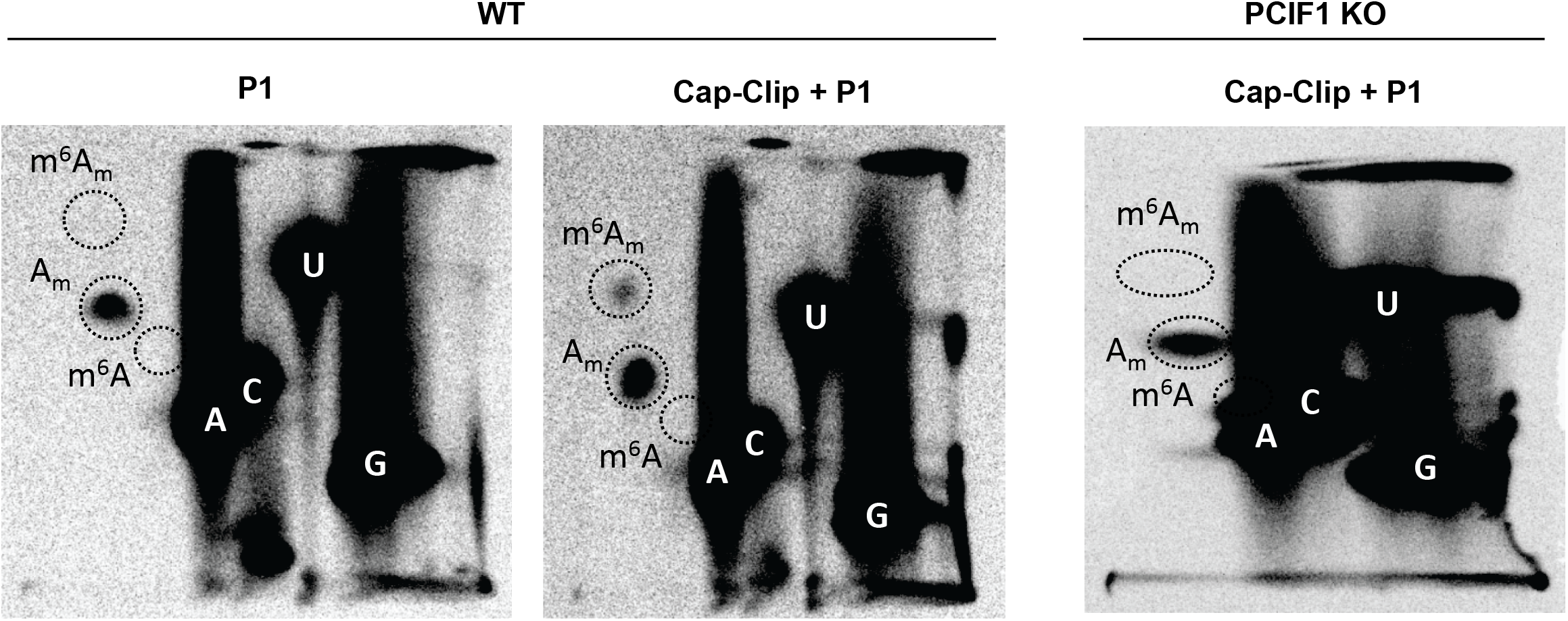
VSV mRNA in A549 cells contain m^6^A_m_. A549 WT or PCIF1 KO cells were infected with VSV at a MOI of 5, and viral mRNA specifically radiolabeled, extracted, and digested as in Fig 2A. Released nucleotide monophosphates were resolved by 2D-TLC and detected by phosphorimaging (representative images, n=3) **(A)** VSV mRNA from wild type cells digested with the indicated enzymes. **(B)** VSV mRNA from PCIF1 KO cells.

**Figure S9:**
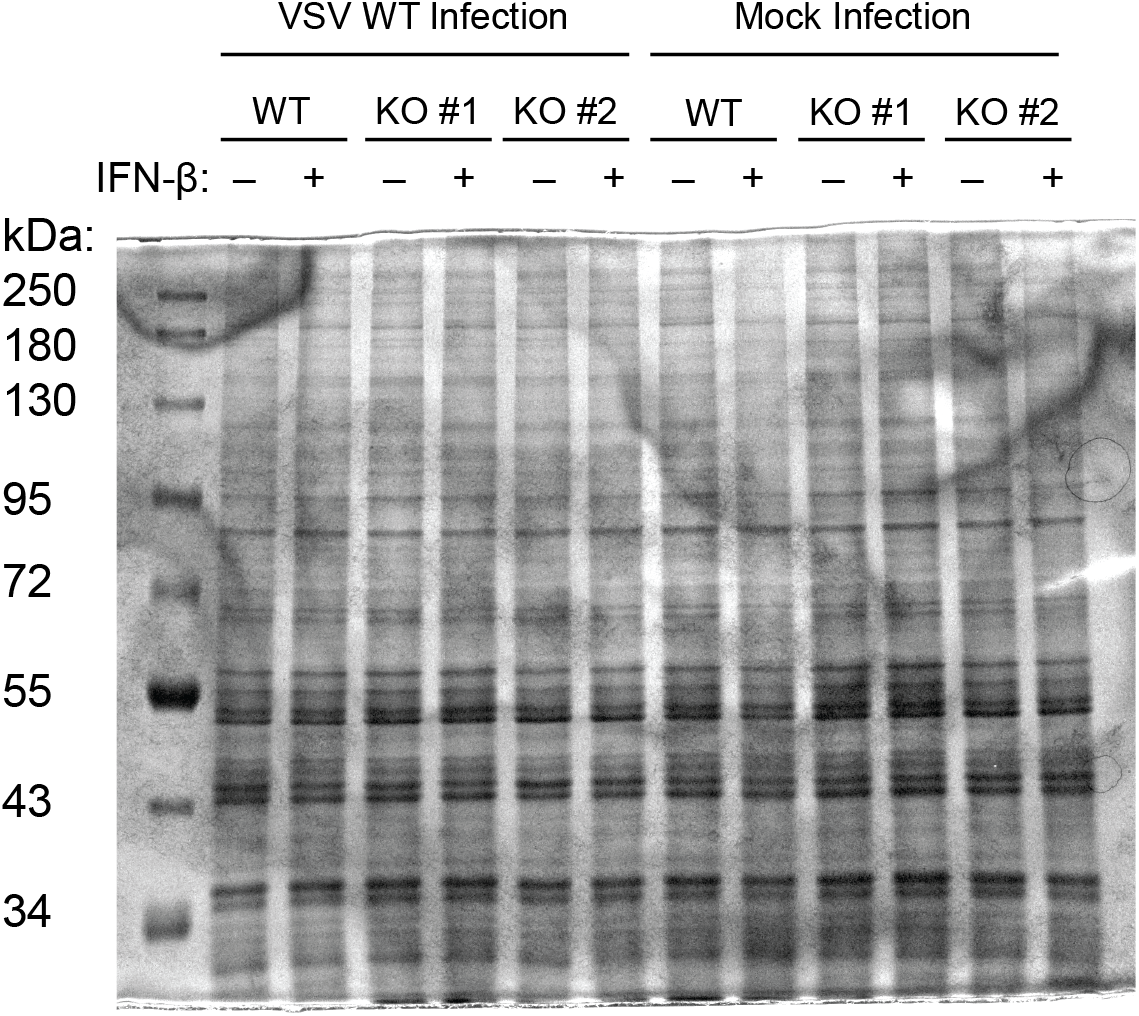
Loading control and protein markers for Fig 5D. Gels used for PAGE-analysis of radiolabeled products in Fig 5D were stained with 0.25% Coomassie Brilliant Blue G-250 in 10% acetic acid, followed by destaining in 10% acetic acid. Gels were visualized by OD laser scanning on a GE Typhoon 5. Staining indicated even protein loading (representative image shown corresponding to autoradiogram in Fig 5D, n=3).

**Table S1:**
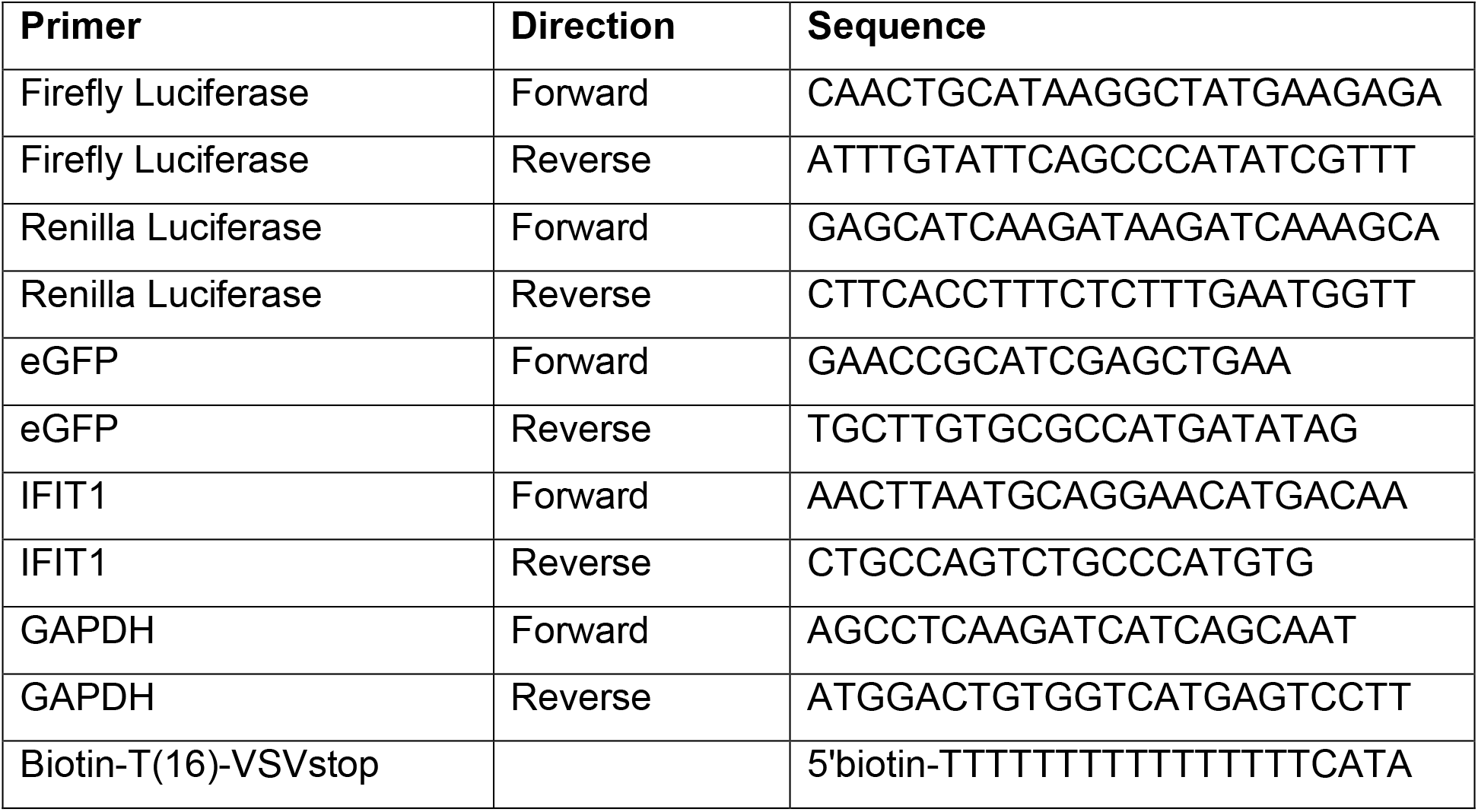
Oligonucleotides used for qPCR or mRNA selection.

## Notes

### Competing Interest Statement

The authors have declared no competing interest.

## References

1. Furuichi Y (2015) Discovery of m(7)G-cap in eukaryotic mRNAs. Proceedings of the Japan Academy. Series B, Physical and biological sciences 91(8):394–409.

2. Furuichi Y, LaFiandra A, & Shatkin AJ (1977) 5’-Terminal structure and mRNA stability. Nature 266(5599):235–239.

3. Furuichi Y & Shatkin AJ (2000) Viral and cellular mRNA capping: past and prospects. Advances in virus research 55:135–184.

4. Leung DW & Amarasinghe GK (2016) When your cap matters: structural insights into self vs non-self recognition of 5’ RNA by immunomodulatory host proteins. Current opinion in structural biology 36:133–141.

5. Shuman S (1995) Capping enzyme in eukaryotic mRNA synthesis. Progress in nucleic acid research and molecular biology 50:101–129.

6. Hornung V, et al. (2006) 5’-Triphosphate RNA is the ligand for RIG-I. Science 314(5801):994–997.

7. Hyde JL & Diamond MS (2015) Innate immune restriction and antagonism of viral RNA lacking 2-O methylation. Virology 479-480:66–74.

8. Meyer KD & Jaffrey SR (2014) The dynamic epitranscriptome: N6-methyladenosine and gene expression control. Nature reviews. Molecular cell biology 15(5):313–326.

9. Meyer KD & Jaffrey SR (2017) Rethinking m(6)A Readers, Writers, and Erasers. Annual review of cell and developmental biology 33:319–342.

10. Keith JM, Ensinger MJ, & Mose B (1978) HeLa cell RNA (2’-O-methyladenosine-N6-)-methyltransferase specific for the capped 5’-end of messenger RNA. The Journal of biological chemistry 253(14):5033–5039.

11. Akichika S, et al. (2019) Cap-specific terminal N (6)-methylation of RNA by an RNA polymerase II-associated methyltransferase. Science 363(6423).

12. Boulias K, et al. (2019) Identification of the m(6)Am Methyltransferase PCIF1 Reveals the Location and Functions of m(6)Am in the Transcriptome. Molecular cell 75(3):631–643 e638.

13. Sendinc E, et al. (2019) PCIF1 Catalyzes m6Am mRNA Methylation to Regulate Gene Expression. Molecular cell 75(3):620–630 e629.

14. Sun H, Zhang M, Li K, Bai D, & Yi C (2019) Cap-specific, terminal N(6)-methylation by a mammalian m(6)Am methyltransferase. Cell research 29(1):80–82.

15. Wei C, Gershowitz A, & Moss B (1975) N6, O2’-dimethyladenosine a novel methylated ribonucleoside next to the 5’ terminal of animal cell and virus mRNAs. Nature 257(5523):251–253.

16. Wei CM, Gershowitz A, & Moss B (1976) 5’-Terminal and internal methylated nucleotide sequences in HeLa cell mRNA. Biochemistry 15(2):397–401.

17. Mauer J, et al. (2017) Reversible methylation of m(6)Am in the 5’ cap controls mRNA stability. Nature 541(7637):371–375.

18. Wei J, et al. (2018) Differential m(6)A, m(6)Am, and m(1)A Demethylation Mediated by FTO in the Cell Nucleus and Cytoplasm. Molecular cell 71(6):973–985 e975.

19. Baltimore D, Huang AS, & Stampfer M (1970) Ribonucleic acid synthesis of vesicular stomatitis virus, II. An RNA polymerase in the virion. Proceedings of the National Academy of Sciences of the United States of America 66(2):572–576.

20. Liang B, et al. (2015) Structure of the L Protein of Vesicular Stomatitis Virus from Electron Cryomicroscopy. Cell 162(2):314–327.

21. Barr JN, Whelan SP, & Wertz GW (1997) cis-Acting signals involved in termination of vesicular stomatitis virus mRNA synthesis include the conserved AUAC and the U7 signal for polyadenylation. Journal of virology 71(11):8718–8725.

22. Stillman EA & Whitt MA (1997) Mutational analyses of the intergenic dinucleotide and the transcriptional start sequence of vesicular stomatitis virus (VSV) define sequences required for efficient termination and initiation of VSV transcripts. Journal of virology 71(3):2127–2137.

23. Wang JT, McElvain LE, & Whelan SP (2007) Vesicular stomatitis virus mRNA capping machinery requires specific cis-acting signals in the RNA. Journal of virology 81(20):11499–11506.

24. Moyer SA & Banerjee AK (1976) In vivo methylation of vesicular stomatitis virus and its host-cell messenger RNA species. Virology 70(2):339–351.

25. Neidermyer WJ, Jr. & Whelan SPJ (2019) Global analysis of polysome-associated mRNA in vesicular stomatitis virus infected cells. PLoS pathogens 15(6):e1007875.

26. Knipe DM & Howley P (2013) Fields Virology (Wolters Kluwer, Lippincott Williams & Wilkins., Philadelphia) 6 Ed.

27. Kruse S, et al. (2011) A novel synthesis and detection method for cap-associated adenosine modifications in mouse mRNA. Scientific reports 1:126.

28. Rahmeh AA, Li J, Kranzusch PJ, & Whelan SP (2009) Ribose 2’-O methylation of the vesicular stomatitis virus mRNA cap precedes and facilitates subsequent guanine-N-7 methylation by the large polymerase protein. Journal of virology 83(21):11043–11050.

29. Li J, Wang JT, & Whelan SP (2006) A unique strategy for mRNA cap methylation used by vesicular stomatitis virus. Proceedings of the National Academy of Sciences of the United States of America 103(22):8493–8498.

30. Wang G, Kouwaki T, Okamoto M, & Oshiumi H (2019) Attenuation of the Innate Immune Response against Viral Infection Due to ZNF598-Promoted Binding of FAT10 to RIG-I. Cell reports 28(8):1961–1970 e1964.

31. Abbas YM, et al. (2017) Structure of human IFIT1 with capped RNA reveals adaptable mRNA binding and mechanisms for sensing N1 and N2 ribose 2’-O methylations. Proceedings of the National Academy of Sciences of the United States of America 114(11):E2106–E2115.

32. Johnson B, et al. (2018) Human IFIT3 Modulates IFIT1 RNA Binding Specificity and Protein Stability. Immunity 48(3):487–499 e485.

33. Li Y, Dai J, Song M, Fitzgerald-Bocarsly P, & Kiledjian M (2012) Dcp2 decapping protein modulates mRNA stability of the critical interferon regulatory factor (IRF) IRF-7. Molecular and cellular biology 32(6):1164–1172.

34. Williams GD, Gokhale NS, Snider DL, & Horner SM (2020) The mRNA Cap 2’-O-Methyltransferase CMTR1 Regulates the Expression of Certain Interferon-Stimulated Genes. mSphere 5(3).

35. Pandey RR, et al. (2020) The Mammalian Cap-Specific m(6)Am RNA Methyltransferase PCIF1 Regulates Transcript Levels in Mouse Tissues. Cell reports 32(7):108038.

36. Limbach PA, Crain PF, & McCloskey JA (1994) Summary: the modified nucleosides of RNA. Nucleic acids research 22(12):2183–2196.

37. You C, Dai X, & Wang Y (2017) Position-dependent effects of regioisomeric methylated adenine and guanine ribonucleosides on translation. Nucleic acids research 45(15):9059–9067.

38. Hudson BH & Zaher HS (2015) O6-Methylguanosine leads to position-dependent effects on ribosome speed and fidelity. Rna 21(9):1648–1659.

39. Sommer S, et al. (1976) The methylation of adenovirus-specific nuclear and cytoplasmic RNA. Nucleic acids research 3(3):749–765.

40. Moss B & Koczot F (1976) Sequence of methylated nucleotides at the 5’-terminus of adenovirus-specific RNA. Journal of virology 17(2):385–392.

41. Haegeman G & Fiers W (1978) Characterization of the 5’-terminal capped structures of late simian virus 40-specific mRNA. Journal of virology 25(3):824–830.

42. Moss B, Gershowitz A, Stringer JR, Holland LE, & Wagner EK (1977) 5’-Terminal and internal methylated nucleosides in herpes simplex virus type 1 mRNA. Journal of virology 23(2):234–239.

43. Flavell AJ, Cowie A, Legon S, & Kamen R (1979) Multiple 5’ terminal cap structures in late polyoma virus RNA. Cell 16(2):357–371.

44. Boone RF & Moss B (1977) Methylated 5’-terminal sequences of vaccinia virus mRNA species made in vivo at early and late times after infection. Virology 79(1):67–80.

45. Faul EJ, Lyles DS, & Schnell MJ (2009) Interferon response and viral evasion by members of the family rhabdoviridae. Viruses 1(3):832–851.

46. Horwitz JA, Jenni S, Harrison SC, & Whelan SPJ (2020) Structure of a rabies virus polymerase complex from electron cryo-microscopy. Proceedings of the National Academy of Sciences of the United States of America 117(4):2099–2107.

47. Ogino M, Ito N, Sugiyama M, & Ogino T (2016) The Rabies Virus L Protein Catalyzes mRNA Capping with GDP Polyribonucleotidyltransferase Activity. Viruses 8(5).

48. Chandran K, Sullivan NJ, Felbor U, Whelan SP, & Cunningham JM (2005) Endosomal proteolysis of the Ebola virus glycoprotein is necessary for infection. Science 308(5728):1643–1645.

49. Cureton DK, Burdeinick-Kerr R, & Whelan SP (2012) Genetic inactivation of COPI coatomer separately inhibits vesicular stomatitis virus entry and gene expression. Journal of virology 86(2):655–666.

50. Cureton DK, Massol RH, Saffarian S, Kirchhausen TL, & Whelan SP (2009) Vesicular stomatitis virus enters cells through vesicles incompletely coated with clathrin that depend upon actin for internalization. PLoS pathogens 5(4):e1000394.

51. Whelan SP, Ball LA, Barr JN, & Wertz GT (1995) Efficient recovery of infectious vesicular stomatitis virus entirely from cDNA clones. Proceedings of the National Academy of Sciences of the United States of America 92(18):8388–8392.

52. Whelan SP & Wertz GW (2002) Transcription and replication initiate at separate sites on the vesicular stomatitis virus genome. Proceedings of the National Academy of Sciences of the United States of America 99(14):9178–9183.

